# ProteasomeID: quantitative mapping of proteasome interactomes and substrates for *in vitro* and *in vivo* studies

**DOI:** 10.1101/2022.08.09.503299

**Authors:** Aleksandar Bartolome, Julia C. Heiby, Domenico Di Fraia, Ivonne Heinze, Hannah Knaudt, Ellen Späth, Omid Omrani, Alberto Minetti, Maleen Hofmann, Joanna M. Kirkpatrick, Therese Dau, Alessandro Ori

**Author notes:** equal contribution (Julia C. Heiby, Domenico Di Fraia).

## Abstract

Proteasomes are essential molecular machines responsible for the degradation of proteins in eukaryotic cells. Altered proteasome activity has been linked to neurodegeneration, auto-immune disorders and cancer. Despite the relevance for human disease and drug development, no method currently exists to monitor proteasome composition and interactions *in vivo* in animal models. To fill this gap, we developed a strategy based on tagging of proteasomes with promiscuous biotin ligases and generated a new mouse model enabling the quantification of proteasome interactions by mass spectrometry. We show that biotin ligases can be incorporated in fully assembled proteasomes without negative impact on their activity. We demonstrate the utility of our method by identifying novel proteasome-interacting proteins, charting interactomes across mouse organs, and showing that proximity-labeling enables the identification of both endogenous and small molecule-induced proteasome substrates.

## INTRODUCTION

The ubiquitin–proteasome system (UPS) is a major selective protein degradation system of eukaryotic cells ^1^. Proteasomes influence crucial processes in the cell including protein homeostasis, DNA repair, signal transduction and immune responses ^2^ by degrading a multitude of regulatory, short-lived, damaged and misfolded proteins ^3**_**4^. Impairment of proteasome function has been described to occur during aging ^5^ and in neurodegenerative diseases ^6^ ^1^, and high expression levels of proteasome subunits have been linked to longevity in different species including worms ^7^, fish ^8^ and flies ^9^ ^10^. While partial proteasome inhibition is sufficient to induce cellular senescence ^11^ ^12^, prolonged inhibition of the proteasome induces apoptosis^13^. This has been exploited in cancer therapies, e.g., against blood cancers, as cancer cells are especially sensitive to proteotoxic stress ^14^. Furthermore, the role of proteasomes extends beyond the degradation of proteins. In the immune system, peptide generation by specialized immunoproteasomes regulates antigen presentation and has been linked to auto-immune disorders ^15^.

The eukaryotic proteasome consists of two subcomplexes: the “core particle“ (20S proteasome) that can be present either alone or in association with one (26S proteasome) or two 19S “regulatory particles“ (30S proteasome). These proteasome variants are considered constitutive and they are generated in most of the cells. In addition to the 19S regulatory particles, the core particle can be associated with different alternative regulatory complexes including the proteasome activator PA200 (named PSME4 in mammals) or three different versions of the 11S regulator complex PA28 – PA28α (PSME1), PA28β (PSME2) and PA28γ (PSME3). These are ATP-independent proteasome regulators that modulate proteasome activity towards small peptides and unfolded proteins and can facilitate protein degradation independent of ubiquitin ^16,17^.

The composition of cellular proteasomes can be heterogeneous due to the assembly of distinct sub-complexes in variable stoichiometries, and to the incorporation of tissue/cell type specific components. Specialized proteasomes have been described in immune cells, thymus, sperm cells ^18^, and cancer ^19^. Sub-populations of proteasomes have been shown to be located in different cellular compartments including inner nuclear membrane ^20^, endoplasmic reticulum ^21^, Golgi apparatus ^22^, plasma membranes ^23^, primary cilia ^24^, and protein aggregates ^25^. Moreover, the assembly state of proteasomes and their biophysical properties can be modulated in response to, e.g., hyperosmotic stress ^26^ or amino acid starvation ^27^.

Due to the dynamic nature of proteasome composition and its distribution within cells, it is important to employ analytical approaches that can capture protein-protein interactions across different cellular compartments and account for transient interactions. Traditional methods such as co-immunoprecipitation, affinity purification or cell fractionation coupled to mass spectrometry have been valuable in studying proteasome interactions ^28–32^, but they may not capture transient or weak interactions. To address this limitation, proximity labeling assays, such as BioID or APEX, utilize engineered proteins that can biotinylate neighboring proteins in close proximity within the cellular context, and thereby enable a more comprehensive and dynamic characterization of protein-protein interactions ^33^.

Here, we present an approach named ProteasomeID that enables the quantification of proteasome interacting proteins and substrates in cultured human cells and mouse models. The approach entails fusing the proteasome with promiscuous biotin ligases, which are incorporated into fully assembled proteasomes without affecting their activity. The ligases label proteins with biotin that come into proximity (∼10 nm) of the tagged proteasome subunit. Biotinylated proteins are then captured from cell or tissue lysates using an optimized streptavidin enrichment protocol and analyzed by deep Data Independent Acquisition (DIA) mass spectrometry. We show that ProteasomeID can quantitatively monitor the majority of proteasome interacting proteins both *in situ* and *in vivo* in mice, and identify novel proteasome interacting proteins. In addition, when combined with proteasome inhibition, ProteasomeID enables the identification of proteasome substrates, including low abundant transcription factors.

## RESULTS

### Design of a proximity labeling strategy to monitor proteasome interactions

First, we tested multiple locations of the biotin ligase to ensure that the integration within the proteasome complex would not impede its assembly or disrupt its functionality. Based on previous studies where proteasome members were tagged with fluorescent proteins in mammalian cells ^34,35^, we fused the promiscuous biotin ligase BirA* either to the C-termini of the 20S core particle protein PSMA4/ɑ3, the 19S particle base protein PSMC2/Rpt1 or the N-terminus of the 19S particle lid protein PSMD3/Rpn3 (Figure 1a and S1a). Each construct also contained a FLAG tag for fusion protein detection. We used these constructs to generate stable HEK293 FlpIn TREx (HEK293T) cell lines that overexpress the BirA* fusion proteins under the control of a tetracycline inducible promoter. A cell line expressing only the BirA* protein was used as a control to account for non specific biotinylation. We confirmed tetracycline-dependent expression of the corresponding cell lines by anti-FLAG immunoblot, and confirmed biotinylating activity following supplementation of exogenous biotin using streptavidin-HRP blot (Figure 1b). These results were validated by immunofluorescence analysis (Figure 1c and S1b).

**Figure 1:**
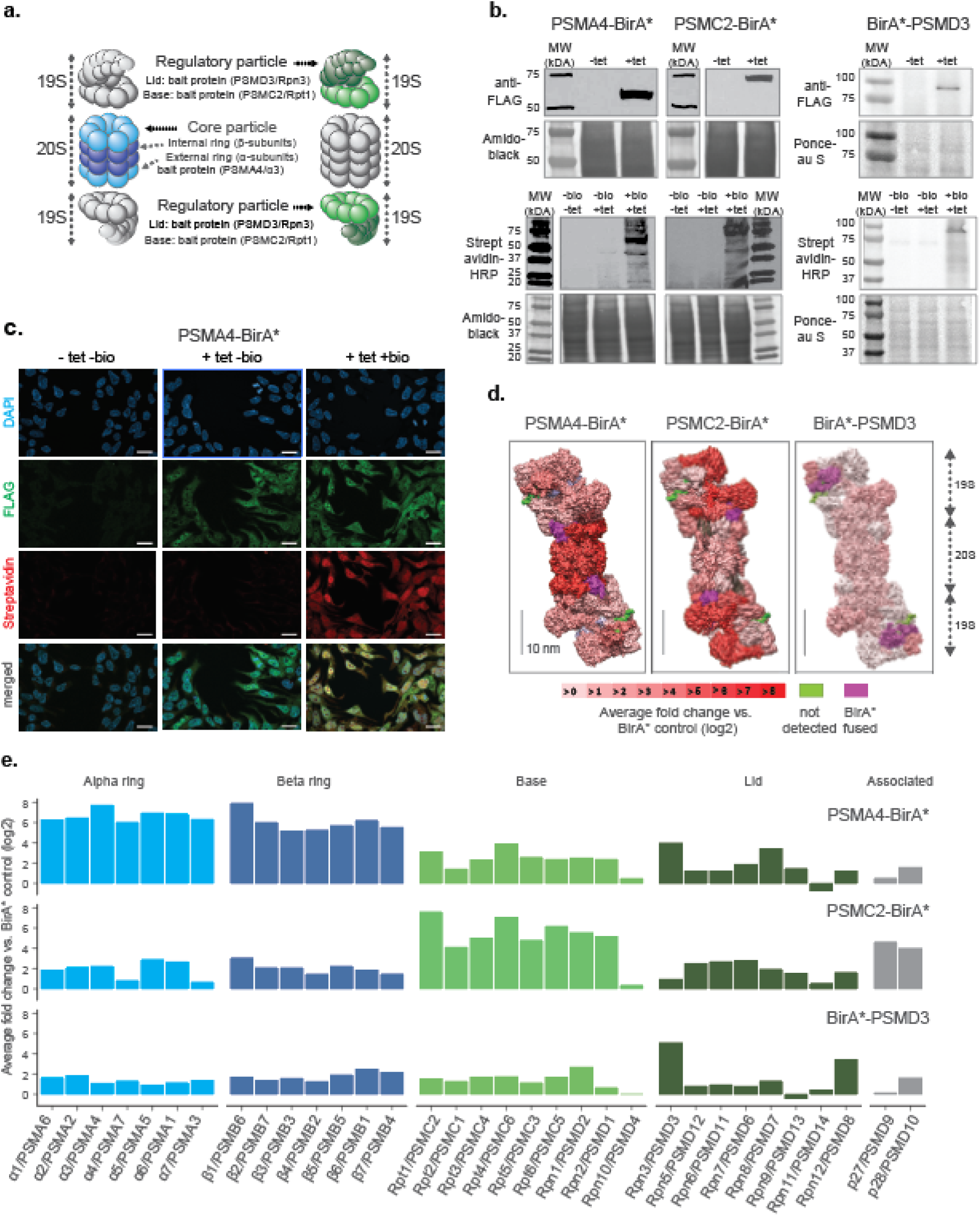
Establishment of a cell culture model system for proximity labeling of proteasomes. a. Schematic representation of proteasome with the substructures containing subunits fused to biotin ligase highlighted in color (shown in 2 shades of blue and green for 20S and 19S proteasome, respectively). b. Upper panel, immunoblot of BirA* fusion proteins performed on lysates from HEK293T cells stably transfected with PSMA4-BirA*-FLAG, PSMC2-BirA*-FLAG or BirA*-FLAG-PSMD3 following 24 h incubation with (+tet) or without (−tet) tetracycline. Lower panel, streptavidin-HRP blot following induction of BirA* fusion proteins with tetracycline and supplementation of biotin for 24 h. Amido Black or Ponceau stainings were used as loading control. HRP: horseradish peroxidase. c. Immunofluorescence analysis of PSMA4-BirA*-FLAG cell line 4 days after seeding without addition of any substance (-tet -bio), with addition of only tetracycline for 4 days (+tet -bio) or with addition of both tetracycline for 4 days and biotin for 1 day (+tet +bio). Scale bar = 20 µm. d. Level of enrichment of proteasome components measured by ProteasomeID in the context of the proteasome structure. Enriched proteins are depicted in different shades of red according to the log2 fold enrichment vs. BirA* control. Scale bar = 10 nm. The proteasome structure depicted was obtained from the PDB:5T0C model of the human 26S proteasome ^62^ and rendered using Chimera ^63^. e. Enrichment level comparison for proteasome components achieved in 3 different cell lines of ProteasomeID. Enriched proteins are depicted in the same color code as in panel a. and according to the log2 fold enrichment vs. BirA* control. n = 4 biological replicates.

In order to identify the most suitable fusion protein for proximity labeling of proteasomes, we compared the enrichment of proteasome components in streptavidin pull downs performed with different cell lines that we generated. We optimized a BioID protocol that we previously developed ^36^ to improve the identification of biotinylated proteins by liquid chromatography tandem mass spectrometry (LC-MS/MS). Briefly, the protocol entails the capture of biotinylated proteins from cell lysates using streptavidin beads followed by enzymatic on-bead digestion and analysis of the resulting peptides by LC-MS/MS. We introduced chemical modification of streptavidin beads and changed the protease digestion strategy to reduce streptavidin contamination following on-bead digestion (Figure S2a). In addition, we improved the data analysis by implementing Data Independent Acquisition (DIA). These optimizations allowed us to drastically reduce (>4 fold) the background from streptavidin-derived peptides (Figure S2b and S2c), and to increase more than 2 fold the number of identified proteins and biotinylated peptides in our BioID experiments (Figure S2d).

Using this optimized BioID protocol, we analyzed samples enriched from cell lines expressing PSMA4-BirA*, PSMC2-BirA*, BirA*-PSMD3 or BirA* control (Figure S2e). We found significant enrichment of proteasome subunits for each of the cell lines expressing BirA* fusion proteins compared to BirA* control (Figure S2f and Table S1). However, the pattern of enrichment varied between fusion proteins (Figure 1d and 1e). PSMA4-BirA* provided the strongest enrichment for 20S proteins, but also a prominent enrichment of other proteasome components (typically >4 fold), while PSMC2-BirA* enriched preferentially 19S base proteins. BirA*-PSMD3 displayed a more homogenous, but less pronounced enrichment of proteasome proteins (typically ∼2 fold). The different enrichment patterns reflect the localization of the fusion proteins within the complex, but it might also indicate interference of the biotin ligase with the assembly of the proteasome, especially in the case of PSMC2-BirA*. Consequently, we decided to focus on the PSMA4-BirA* fusion protein for further characterization as it showed an overall stronger enrichment of proteasomal proteins from both the 20S and 19S particles.

### PSMA4-BirA* is incorporated into fully assembled proteasomes and does not interfere with their proteolytic activity

Next, we performed a series of experiments to confirm the incorporation of PSMA4-BirA* into fully assembled proteasomes and exclude any potential interference with their proteolytic activity. Since the BirA* fusion protein is over-expressed using a CMV promoter, we first confirmed that the abundance levels of PSMA4-BirA* are comparable to the ones of the endogenous PSMA4 using an anti-PSMA4 immunoblot (Figure 2a and S3a).

**Figure 2:**
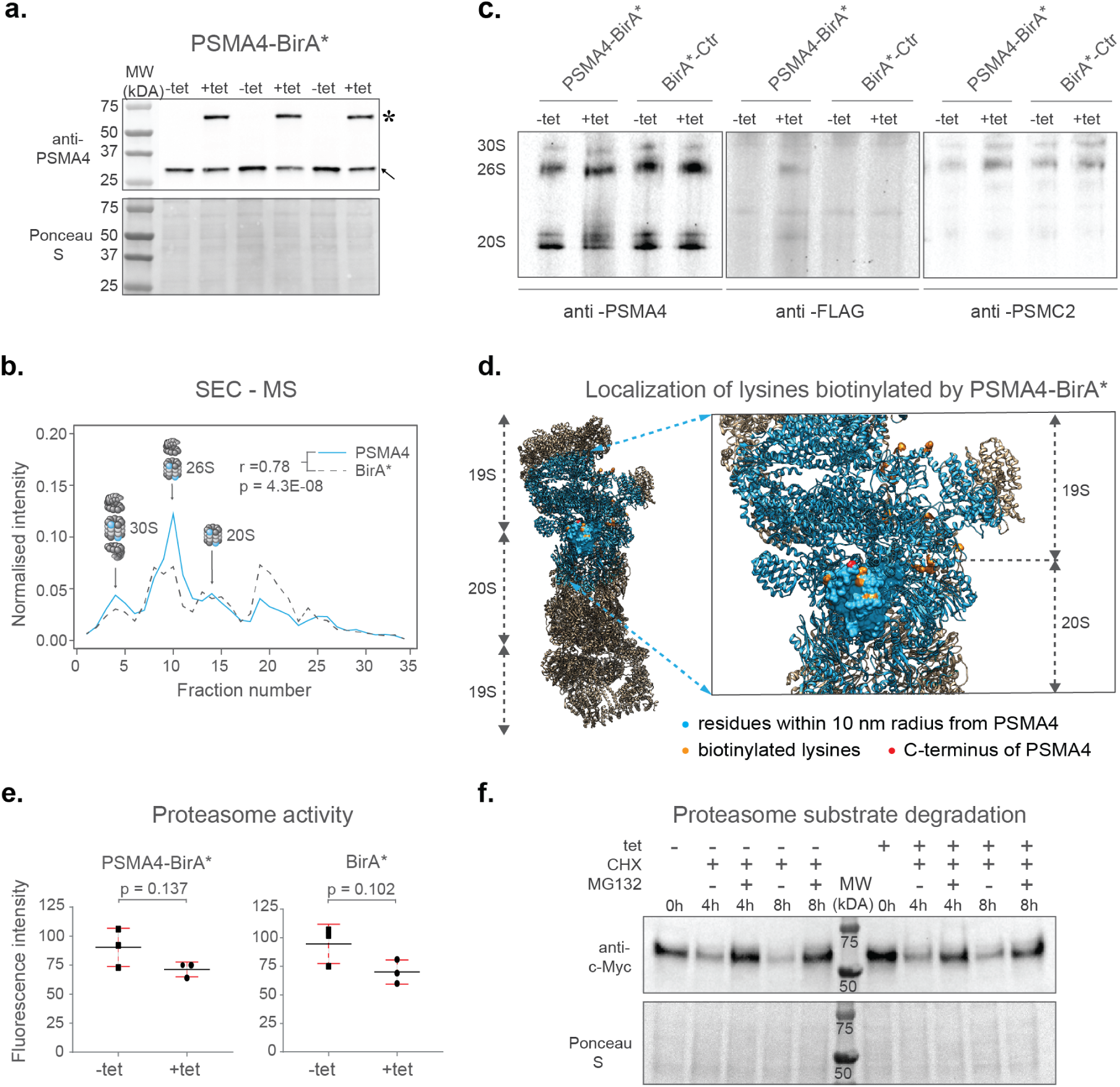
Validation of ProteasomeID. a. Comparison of expression levels of PSMA4-BirA* (lanes marked by star) and its endogenous counterpart (lanes marked by arrowhead), following 24 h incubation with (+tet) or without (−tet) tetracycline. Ponceau S staining was used as loading control. b. Size exclusion chromatography (SEC) analysis of lysates from HEK293T cells stably expressing PSMA4-BirA* following 24 h incubation with tetracycline. SEC fractions were analyzed by DIA mass spectrometry and elution profiles were built for each protein using protein quantity values normalized to the sum of quantities across all fractions. Depicted are elution profiles of PSMA4 (proteasome subunit, blue) and BirA* (biotinylating enzyme, dashed line). The peaks corresponding to different proteasome assemblies were assigned based on the elution profiles of other proteasome components. c. Immunoblot for proteasome subunits PSMA4 (left panel), PSMC2 (right panel) and FLAG tag (middle panel) of cell lysates separated by native PAGE from PSMA4-BirA* and BirA*-Ctr cell lines with and without tetracycline addition. Tet = tetracycline, 30S = indicates position of proteasome structures containing one core and 2 regulatory particles, 26S = indicates position of proteasome structures containing one core and 1 regulatory particle, 20S = indicates position of proteasome structures consisting of only single core particle. d. Biotinylated lysines identified by ProteasomeID. All the residues within a 10 nm radius of the PSMA4 C-terminus are highlighted in cyan. Red color indicates the C-terminus of PSMA4 where BirA* is fused (not present in the structure), and the identified biotinylated lysines are depicted in orange. Only the structure of the modified subunit is depicted with a surface model and all the other subunits are depicted as helix-loop structures. Biotinylated residues were obtained from the ACN fraction of PSMA4-BirA*. The proteasome structure depicted was obtained from the PDB:5T0C model of the human 26S proteasome ^62^ and rendered using Chimera ^63^. e. Proteasome activity assay performed on lysates from cell lines expressing different BirA* fusion proteins, following 24 h incubation with (+tet) or without (−tet) tetracycline. Equal amounts of protein extracts were incubated with proteasome substrate LLVY-7-Amino-4-methylcoumarin (AMC) and substrate cleavage assessed by fluorimetry. n = 3 biological replicates, error bars indicate standard deviation of the mean, paired t-test. f. Cycloheximide-chase experiment on c-Myc stability. PSMA4-BirA*cells were incubated with 50 μg/ml cycloheximide (CHX) for the indicated times in the presence or absence of MG132 (20 μM) and tetracycline (1 µg/µl). Cells lysates were then prepared for Western blot analysis of steady-state levels of c-Myc.Tet = tetracycline, CHX = cycloheximide.

In order to assess the assembly of PSMA4-BirA* into proteasome complexes, we performed size exclusion chromatography coupled to quantitative mass spectrometry (SEC-MS) analysis of the cell line expressing PSMA4-BirA* following induction by tetracycline. Protein elution profiles built from mass spectrometry data obtained from 32 SEC fractions revealed three major distinct peaks corresponding to the major assembly states of the proteasome (Table S2). These include 30S proteasomes, containing a core particle capped with 2 regulatory particles, 26S proteasomes, containing a core particle capped with 1 regulatory particle, and isolated core particles (20S proteasomes) (Figure 2b and S3b). With the exception of a peak in lower molecular weight fractions (∼fraction 20), likely representing intermediate complex assemblies, the majority of the BirA* signal (∼60%) correlated with the elution profile of other proteasome components in all the assembly states (Figure S3b). We have additionally performed anti-FLAG immunoblot upon blue native gel electrophoresis (BN-PAGE) to confirm the incorporation of PSMA4-BirA* into assembled proteasomes, and to demonstrate that the assembly of proteasomes is not affected by the overexpression of PSMA4-BirA* (Figure 2c). Finally, we used our mass spectrometry data to identify sites of protein biotinylation in PSMA4-BirA* expressing cells. By mapping the identified sites on the proteasome structure, we could confirm the specificity of protein biotinylation by showing that 25 out of 26 residues identified as biotinylated by PSMA4-BirA* are located less than 10 nm away from the C-terminus of PSMA4 (Figure 2d and Table S2). Importantly, 18 of these sites were located on proteins from the 19S regulatory particle, further supporting the incorporation of PSMA4-BirA* into assembled proteasomes.

In order to test the influence of BirA* fusion proteins on proteasome function, we measured proteasome chymotrypsin-like activity in cell lysates from BirA* expressing cell lines in presence or absence of tetracycline. We observed a slight, not-significant reduction of proteasome activity (∼15-20%) following addition of tetracycline that was comparable between cell lines expressing proteasome PSMA4-BirA* or BirA* control (Figure 2e). In addition, we tested the impact of PSMA4-BirA* overexpression on the degradation of c-Myc, a known proteasome substrate, using a cycloheximide chase experiment. We showed that tetracycline mediated induction of PSMA4-BirA* did not have a major effect on the degradation kinetics of c-Myc (Figure 2f and S3c). Also, levels of K48-ubiquitylated proteins were not affected (Figure S3d), further confirming that PSMA4-BirA* has no major impact on the proteolytic activity of the proteasome in cells.

### ProteasomeID retrieves proteasome subunits, assembly factors and known proteasome interactors

To assess the ability of ProteasomeID to retrieve known proteasome interacting proteins (PIPs) in addition to proteasome components, we implemented a logistic regression classifier algorithm (Figure 3a). The classifier is based on an “Enrichment score” that we obtained by combining the average log2 ratio and the negative logarithm of the q value from differential protein abundance analysis performed vs. BirA* control line. We then used a set of true positives, here, proteasome members, and true negatives, here, mitochondrial matrix proteins, which should not interact directly with the proteasome under homeostatic conditions, to identify a cut-off for defining ProteasomeID enriched proteins at a controlled false positive rate (Figure 3b). We validated the classifier using a receiver operator characteristic (ROC) analysis (Figure S3e). Using the classifier, we identified 608 protein groups enriched by ProteasomeID for the PSMA4-BirA* dataset (false positive rate, FPR < 0.05, Figure 3c and Table S3).

**Figure 3:**
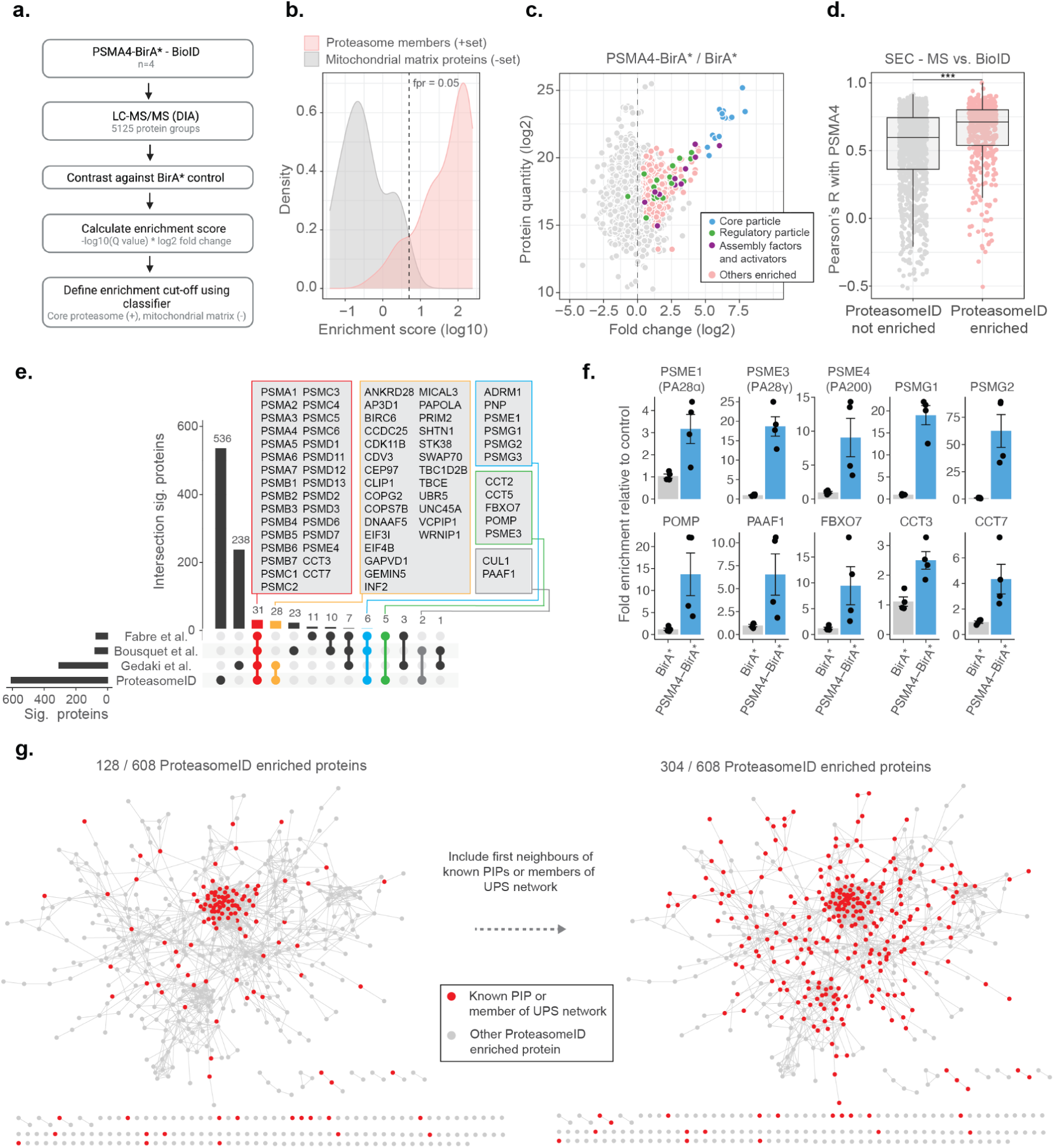
ProteasomeID identifies known proteasome interactors. a. Schematic depiction of the classifier algorithm used to unbiasedly define proteins enriched by ProteasomeID. The classifier is based on an “Enrichment score” obtained by combining the average log2 ratio and the negative logarithm of the q value from differential protein abundance analysis performed vs. BirA* control line. b. Distribution of enrichment scores calculated by the classifier algorithm for proteasome subunits (set of true positives) and mitochondrial matrix proteins (set of true negatives). The dashed vertical line indicates the enrichment score cut-off to define ProteasomeID enriched proteins at FPR < 0.05. c. MA plot of proteins enriched by streptavidin pull-down and analyzed by DIA mass spectrometry. Highlighted in color are proteasome members, assembly factors and activators, and other proteins significantly enriched in ProteasomeID (FPR < 0.05). n = 4 biological replicates. d. Comparison of co-elution profiles obtained by SEC-MS and proteins enriched in ProteasomeID. Pearson correlation values were calculated between PSMA4 and all the other proteins quantified in SEC-MS (n = 4680). Correlation values were compared between proteins significantly enriched in PSMA4-BirA* vs. BirA* and all the other proteins quantified in the ProteasomeID experiment. *** p < 0.001 Wilcoxon Rank Sum test with continuity correction. e. Upset plot showing overlap between ProteasomeID enriched proteins and previous studies that investigated proteasome interacting proteins (PIPs). Different subsets of overlapping proteins are highlighted in color framed boxes. f. Bar plots comparing the levels of enrichment obtained in ProteasomeID experiment for proteasome activators, assembly factors and known PIPs. Enrichment levels were normalized to the levels detected in BirA* control cell line which was set to 1. Protein quantities were derived from DIA mass spectrometry data. Data are shown as mean ± standard error from n = 4 biological replicates. g. Network analysis of 608 interactors of PSMA4-BirA* obtained by ProteasomeID. Identified proteins were filtered for significance by a cutoff of log2 fold change > 1 and Q value < 0.05 in relation to BirA* control. Nodes representing identified proteins that are known PIPs or members of ubiquitin proteasome system (UPS) (left network) or identified protein and their first interacting neighbor are known PIPs or members of UPS (right network) are highlighted in red color. Edges represent high confidence (>0.7) protein-protein interactions derived from the STRING database ^37^.

We investigated the occurrence of PIPs among ProteasomeID enriched proteins using the two approaches. First, we compared the BioID data from PSMA4-BirA* to the SEC-MS data obtained from the same cell line. We could show that SEC elution profiles of proteins enriched in ProteasomeID tend to have significantly higher correlation with the elution profile of PSMA4 (Figure 3d). Since correlated SEC elution profiles are typically interpreted as an indication of physical association in this type of experiment, these data suggest that ProteasomeID can enrich PIPs. Second, we compared our dataset to these three previous proteomic studies that investigate PIPs using complementary approaches: Fabre et al. ^29^ used fractionation by glycerol gradient ultracentrifugation, Bousquet-Dubouch et al. ^30^ used immunoaffinity purification, and Geladaki et al. ^31^ applied cellular fractionation by differential centrifugation. We found 72 proteins overlapping to our dataset including almost all known proteasome members, assembly factors, and activators (Figure 3e and 3f). The overlapping proteins also include members of other complexes known to interact with the proteasome, i.e., the SKP1-CUL1-F-box protein (SCF) complex (CUL1 and FBOX7), and the chaperonin containing TCP-1 (CCT) complex (CCT2, CCT3, CCT5 and CCT7). Some proteins common to at least two of previous studies but were missing in our dataset include immunoproteasome subunits (PSMB10/LMP10, PSMB8/LMP7) that are not expressed in HEK293 cells, the shuttling factor RAD23B and ubiquitin-related proteins including USP14, UCHL5 and UBE3C. Manual inspection of proteins identified exclusively by ProteasomeID revealed additional 37 proteins that were reported in additional studies that investigated proteasome interactions (summarized in Table S3), and 19 proteins that have been associated to the ubiquitin proteasome system (UPS) (Figure S3f). In addition, another 176 proteins found in the ProteasomeID network are interactors of previously reported proteasome interacting proteins, according to high confidence interactions from the STRING database ^37^ (Figure 3g). Therefore, in total, 304 out of 608 (50%) ProteasomeID enriched proteins could be matched to known proteasome-interacting proteins or their direct binding partners. The remaining 304 proteins, which are enriched in GO terms related to microtubule and cilia organization (Table S3), likely represent novel interacting proteins, or proteins that come into proximity of the proteasome without physically interacting with it, or false positives due to the presence of a subpopulation of unassembled PSMA4-BirA*. Comparing the enrichment scores, however, suggest that they are likely proteins that come into proximity of the proteasome without physically interacting with it (Figure S3g).

### Identification of proteasome substrates by ProteasomeID

Having demonstrated that ProteasomeID can be used to obtain snapshots of the proteasome-proximal proteome, we next investigated whether we could use this approach to identify proteasome substrates. We reasoned that under steady state conditions the interaction between proteasomes and their substrates might be too short lived to enable efficient biotinylation. Furthermore, the proteolytic cleavage of substrates by the proteasome would eventually make biotinylated peptides impossible to be identified by our standard proteomic workflow based on tryptic peptides. Therefore, we generated HEK293T cell lines expressing PSMA4-miniTurbo or miniTurbo alone, enabling shorter biotinylation time thanks to the enhanced activity of miniTurbo as compared to BirA* ^38^ (Figure S4a and S4b). We tested biotinylation by PSMA4-miniTurbo using immunoblot and confirmed that 2 h were sufficient to achieve biotinylation levels comparable to the one PSMA4-BirA* after 24 h of biotin supplementation (Figure S4c). Furthermore, we confirmed enrichment of proteasome members and interacting proteins (Figure S4d), and observed positive correlation between the enrichments measured using PSMA4-miniTurbo and PSMA4-BirA*, relatively to their respective control lines (Figure S4e). Applying the classifier algorithm, we identified 168 protein groups enriched for PSMA4-miniTurbo (FPR < 0.05, Figure S4f and g).

Next, we included a step of acute inhibition of the proteasome by the potent cell-permeable inhibitor MG132 for 4h (Figure 4a). We confirmed proteasome inhibition by the accumulation of ubiquitylated proteins assessed by immunoblot (Figure S4h). Principal component analysis of the mass spectrometry data obtained from streptavidin enriched proteins revealed a clear separation between samples treated with MG132 or vehicle control (Figure S4i). Comparison of ProteasomeID enriched proteins from cells treated with proteasome inhibitor versus vehicle control revealed a subset of 141 proteins enriched exclusively upon MG132 treatment (Figure 4b and 4c). These include proteasome activators (PSME1/PA28ɑ, PSME2/PA28β, PSME3/ PA28γ) and ubiquitin (Figure 4d and Table S4), consistent with the recruitment of proteasome activators and direct ubiquitylation of proteasome members following inhibition ^39^. Of the remaining 133 proteins, 77 (58%) have been shown in a previous study ^40^ to display increased ubiquitylation (as assessed by GlyGly enrichment and MS analysis) upon proteasome inhibition, and 39 are reported proteasome substrates such as MYC, ATF4 and PINK1 (Figure 4c and 4d and Table S4). In line with the observation that this subset of proteins is enriched for proteasome substrates, their levels were found to increase relatively to the rest of the proteome upon treatment with MG132 (Figure 4e). Notably, when examining the enrichment of these proteins using ProteasomeID, more pronounced effect sizes were observed (Figure 4e). This suggests that the increased enrichment of these proteins cannot be solely attributed to elevated cellular levels, indicating a specific targeting of these proteins to the proteasome upon MG132.

**Figure 4:**
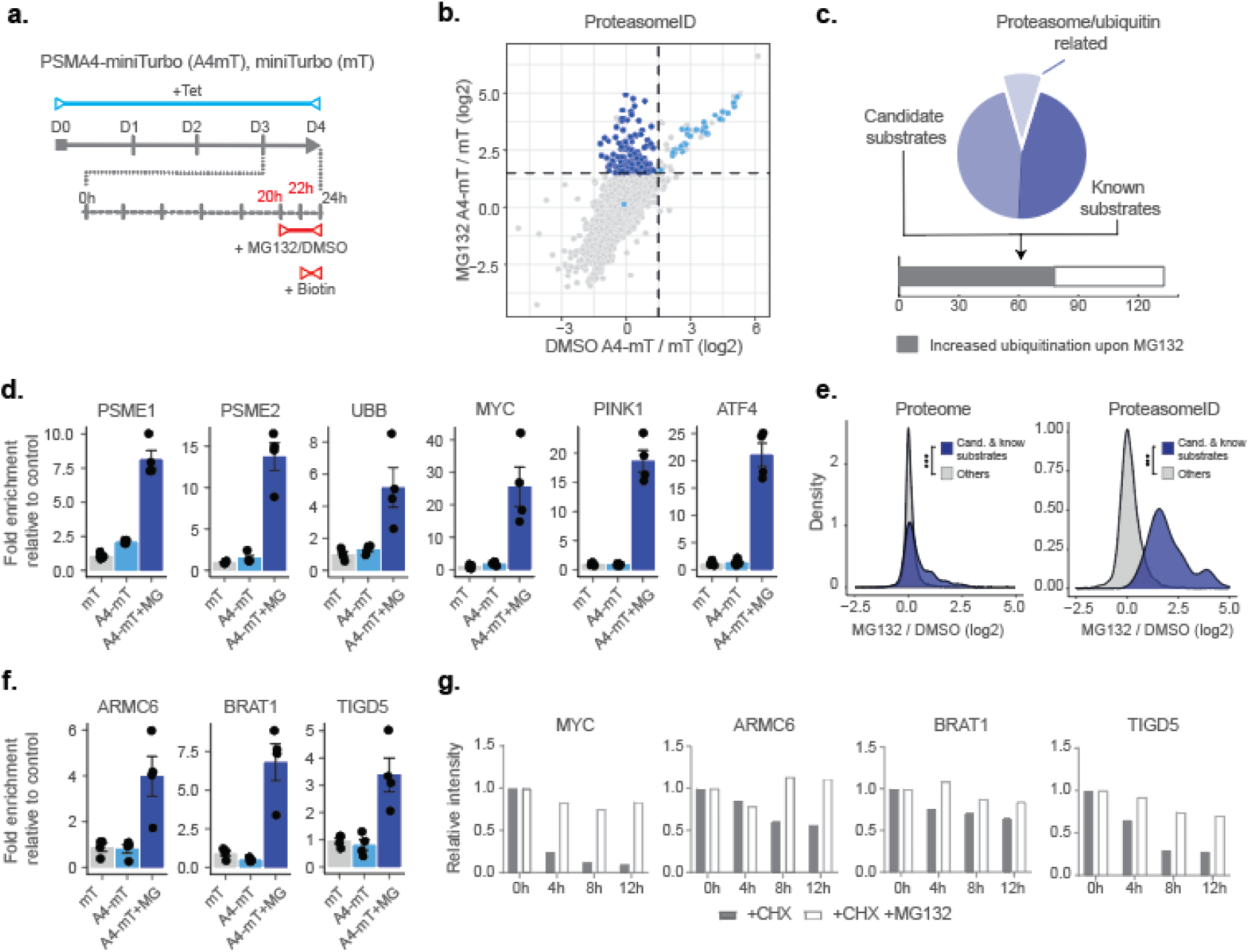
ProteasomeID identifies known and novel endogenous proteasome substrates. a. Scheme of ProteasomeID workflow in HEK293T cells including proteasome inhibition by MG132. PSMA4-miniTurbo expression and incorporation into proteasomes is achieved by 4 day induction with tetracycline. Proteasome inhibition is achieved by addition of 20 µM MG132 4 h before cell harvesting. Biotin substrate for miniTurbo is supplied 2 h before cell harvesting. D: day; h: hour; Tet: tetracycline; Bio: biotin. b. Enriched proteins from ProteasomeID cells treated with proteasome inhibitor MG132 compared to vehicle control. The subset of proteins enriched exclusively upon MG132 treatment are highlighted in dark blue. Data were obtained from n = 4 biological replicates. c. Profile of proteins exclusively enriched upon MG132 treatment of ProteasomeID cells (pie chart). Identified proteins are represented by proteasome/ubiquitin related proteins, known proteasome substrates and potential previously unidentified substrates. The number of proteins identified by this approach for which previous studies showed increased ubiquitylation upon proteasome inhibition is shown in the lower bar plot. d. Bar plots comparing the levels of proteasome activators and ubiquitin, and known proteasome substrates following streptavidin enrichment from different cell lines and following proteasome inhibition by MG132. Protein quantities were derived from DIA mass spectrometry data. mT: miniTurbo control cell line; A4-mT: PSMA4-miniTurbo cell line; I: proteasome inhibition by MG132. Data are shown as mean ± standard error from n = 4 biological replicates. e. Distribution of log2 fold changes following MG132 treatment for candidate and known proteasome substrates identified by ProteasomeID. The fold changes are compared to the other identified proteins using total proteome (left) or ProteasomeID data (right). *** p < 0.001 Wilcoxon Rank Sum test with continuity correction. f. Bar plots comparing the levels of 3 potential novel proteasome substrate proteins following streptavidin enrichment from different cell lines and following proteasome inhibition by MG132. Protein quantities were derived from DIA mass spectrometry data. mT: miniTurbo control cell line; A4-mT: PSMA4-miniTurbo cell line; I: proteasome inhibition by MG132. Data are shown as mean ± standard error from n = 4 biological replicates. g. Cycloheximide-chase experiment on stability of 3 potential novel proteasome substrate proteins. HEK293T cells were incubated with 50 μg/ml cycloheximide (CHX) for the indicated times in the presence or absence of MG132 (20 μM). Cell lysates were then prepared for Western blot analysis of steady-state levels of c-Myc, ARMC6, BRAT1 and TIGD5. c-Myc was used as a positive control as it is a well known proteasome substrate. Densitometric quantification of the bands from the assay are shown in bar plots. For quantification, bands were first normalized to GAPDH as a loading control and subsequently normalized to zero hour, untreated samples (set to 1). CHX = cycloheximide.

In order to validate our findings, we selected 3 proteins exclusively enriched in ProteasomeID upon MG132 that were not previously reported as proteasome substrates (Figure 4f): the BRCA1-associated ATM activator 1 (BRAT1 also known as BAAT1) the armadillo containing protein 6 (ARMC6) and Tigger transposable element-derived protein 5 (TIGD5). While the function of ARMC6 is unknown, BRAT1 has been associated with neurodevelopmental and neurodegenerative disorders ^41^ and shown to play a role in DNA damage response ^42^ and RNA processing ^43^. TIGD5 encodes protein with DNA-binding features. Though its exact function is not established, it has been suggested to operate as a tumor suppressor in ovarian cancer ^44^. We then performed a cycloheximide chase experiment and could validate them as proteasome substrates (Figure 4g and S4j).

Finally, we investigated whether we could use ProteasomeID to identify selective induction of protein degradation by small molecules. For this purpose, we used KB02-JQ1, a well-characterized PROTAC that targets bromodomain-containing proteins (BRDs) for proteasomal degradation ^45^. We pre-treated cells with KB02-JQ1 for 8 hours prior to proteasome inhibition and biotin supplementation (Figure 5a). Mass spectrometry analysis showed global changes induced by KB02-JQ1 either in presence or absence of MG132, as indicated by PCA (Figure S4k). Importantly, we could detect prominent enrichment of BRD containing proteins following treatment with KB02-JQ1 (Figure 5b and Table S4). The effect was more pronounced for BRD2 and BRD3 and less striking for BRD4 (Figure 5c), presumably reflecting different kinetics of induced degradation by KB02-JQ1 ^45^. Notably, the enrichment of BRD2 and BRD3 was also detectable in absence of MG132, in contrast to endogenous substrates that become enriched only following proteasome inhibition (Figure 5c). Together, these data demonstrate that ProteasomeID can be used to detect both endogenous and protein degrader-induced substrates of the proteasome in cultured cells.

**Figure 5:**
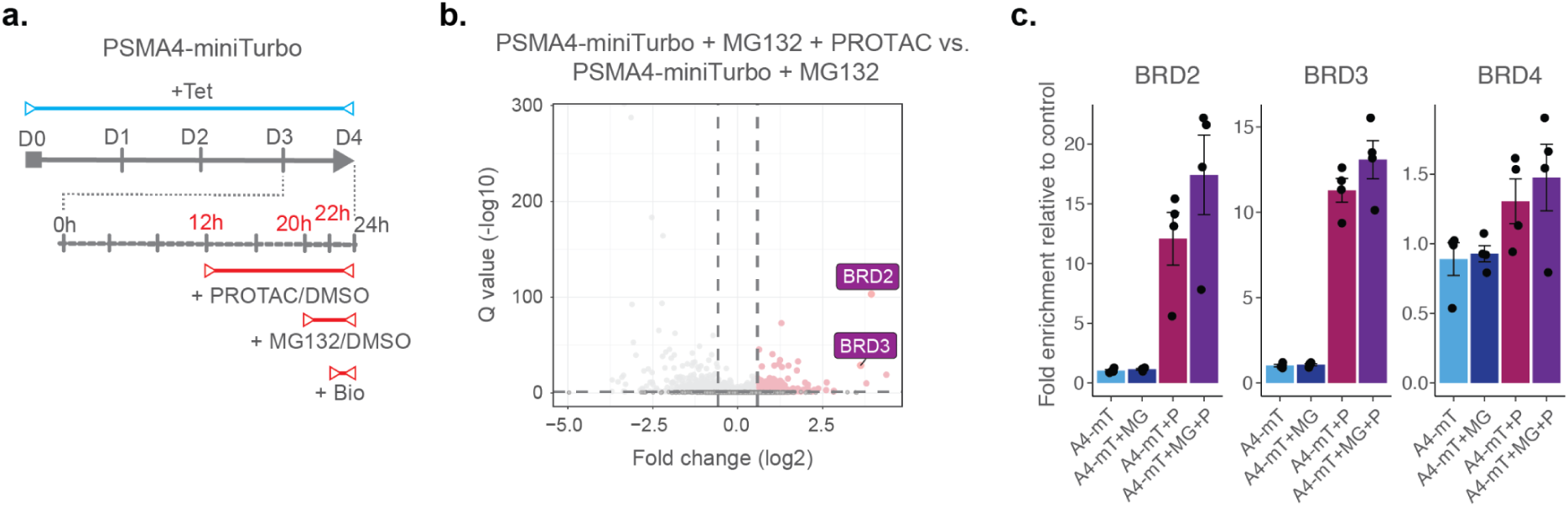
ProteasomeID identifies PROTAC-induced proteasome substrates. a. Scheme of ProteasomeID workflow in HEK293T cells including proteasome inhibition by MG132 and treatment with PROTAC KB02-JQ1. The experimental design is analogous to the one depicted in (Figure 4A) with the additional PROTAC treatment achieved by addition of 10 µM KB02-JQ1 12 h before cell harvesting. D: day; h: hour; Tet: tetracycline; Bio: biotin. b. Volcano plot of proteins enriched by streptavidin pull-down and analyzed by DIA mass spectrometry from PSMA4-miniTurbo cells treated with KB02-JQ1 PROTAC molecule (P) and PSMA4-miniTurbo cells treated with both PROTAC molecule (P) and MG132 proteasome inhibitor (I). Cut offs for enriched proteins: log2 fold change > 1 and Q value < 0.05. n = 4, biological replicates. Enrichment of BRD containing proteins is highlighted in violet boxes. c. Bar plots comparing the levels of BRD-containing proteins following streptavidin enrichment from PSMA4-miniTurbo expressing cells exposed to the proteasome inhibitor MG132 and/or the PROTAC KB02-JQ1. mT: miniTurbo control cell line; A4-mT: PSMA4-miniTurbo cell line; I: proteasome inhibition by MG132; P: PROTAC (KB02-JQ1). Data are shown as mean ± standard error from n = 4 biological replicates.

### A mouse model for *in vivo* ProteasomeID

Having established and validated proximity labeling of proteasomes in a cell culture model, we designed a strategy to implement ProteasomeID in a mouse model (Figure 6a). The mouse model was designed to express the 20S proteasome core particle PSMA4 fused to the biotinylating enzyme miniTurbo and a FLAG tag for the detection of the fusion protein. We chose miniTurbo instead of BirA* because of its higher biotinylating efficiency ^38^. PSMA4-miniTurbo was inserted in the Col1a1 locus downstream of a tetracycline responsive element (TRE) in the D34 mouse embryonic stem cell line ^46^. This line carries a cassette encoding the rtTA3 transactivator and the fluorescent protein mKate on the Rosa 26 locus under the control of a CAG promoter. Importantly, a LoxP-stop-LoxP cassette is present between the CAG promoter and the rtTA3 and mKate expressing cassette, enabling tissue-specific expression via crossing to specific CRE lines. The engineered D34 line was used to generate a mouse line via blastocyst injection. For proof of concept, we crossed the TRE-Psma4-miniTurbo;Rosa26-CAGs-RIK line with a CMV-Cre line that expresses constitutively the CRE recombinase in all tissues ^47^. The obtained TRE-Psma4-miniTurbo;Rosa26-CAGs-RIK line was back crossed to C57BL6/J to remove the CMV-Cre allele. The obtained mouse line constitutively expresses the rtTA3 transactivator, thereby enabling doxycycline inducible expression of the PSMA4-miniTurbo construct in all tissues.

**Figure 6:**
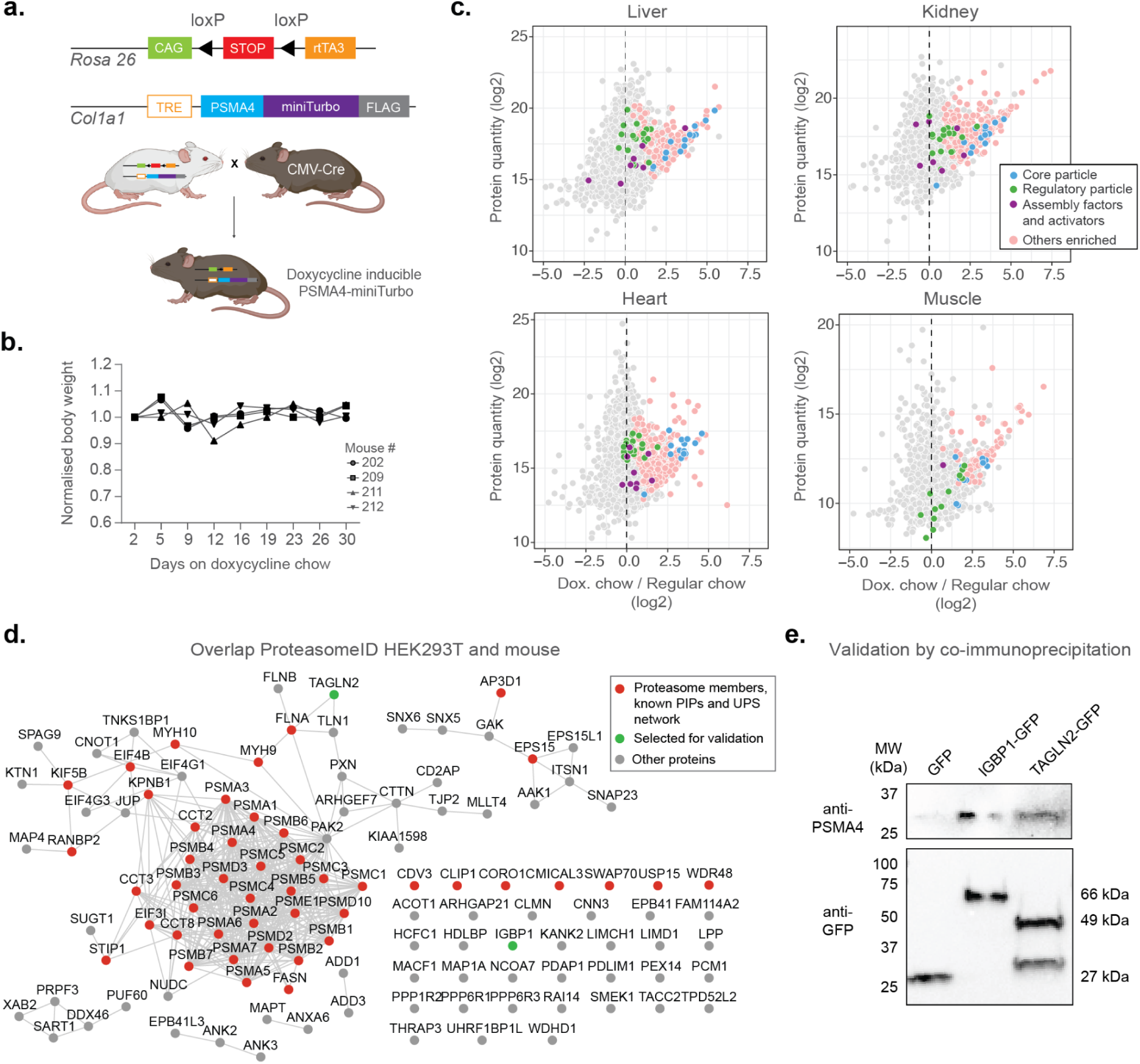
Establishment of a mouse model for *in vivo* ProteasomeID. a. Design of a mouse model for ProteasomeID. The lox-STOP-lox cassette was excised from the Rosa26 locus by crossing with a mouse line expressing the Cre recombinase under the control of an ubiquitous CMV promoter ^47^. CAG: CAG promoter ^64^, TRE: tetracycline-regulated element; rtTA3: reverse tetracycline-dependent transactivator A3 ^46^. (Figure created with biorender). b. Bodyweight curves of the experimental animals. The body weight for each mouse was normalized to its value at day 1 of the experiment (set to 1). c. MA plots of proteins enriched by streptavidin pull-down and analyzed by DIA mass spectrometry from different mouse organs. Highlighted in color are proteasome members, assembly factors and activators, and other proteins significantly enriched in ProteasomeID (FPR < 0.05). n = 4 mice per experimental group. d. Network analysis of overlap of significantly enriched proteins in ProteasomeID from HEK293T cells expressing PSMA4-BirA* and mouse organs (significant in at least one organ). Nodes colored in red indicate proteasome members, PIPs or proteins belonging to the UPS networks. The proteins selected for validation by co-immunoprecipitation are highlighted in green. Edges represent high confidence (>0.7) protein-protein interactions derived from the STRING database ^37^. e. Cells expressing either GFP, IGBP1-GFP or TAGLN2-GFP were used for co-immunoprecipitation using GFP-trap. The elutions from GFP-trap were analyzed by immunoblot using antibodies against PSMA4 or GFP.

We evaluated the induction of PSMA4-miniTurbo in different organs (kidney, liver, heart, skeletal muscle and brain) by feeding mice with doxycycline containing food for 10 or 31 days. We observed no significant changes in body weight nor any sign of suffering (Figure 6b). Immunoblot analysis confirmed expression of PSMA4-miniTurbo in all organs except the brain (Figure S5a). The locus used for PSMA4-miniTurbo expression (Col1a1) is known to be active in all the organs tested ^48^, however the limited penetration of food-administered doxycycline ^49^ limits the usability of our approach in the brain. The temporal dynamics of induction varied between organs. While 10 days of induction were sufficient to achieve PSMA4-miniTurbo protein levels comparable to endogenous PSMA4 in the kidney, other organs required 31 days. In most organs, the expression levels of PSMA4-miniTurbo were comparable to the one of endogenous PSMA4, with the exception of the skeletal muscle. In this organ, the levels of PSMA4-miniTurbo exceeded the endogenous PSMA4, suggesting the existence of a more prominent pool of not incorporated PSMA4-miniTurbo. Based on these observations, we decided to proceed with 31 days of induction and confirmed expression of PSMA4-miniTurbo in the tested organs using anti-FLAG immunohistochemistry (Figure S5b).

*In vivo* BioID requires supplementation of exogenous biotin. We opted for subcutaneous biotin injection based on previous studies ^50^ ^51^ ^52^, and compared 3 x with 7 x daily biotin injections by assessing the recovery of proteasome subunits by LC-MS/MS upon streptavidin enrichment from liver and kidney lysates. We did not observe major quantitative differences in the enrichment of proteasome subunits relatively to control lysates from organs of mice that were not fed with doxycycline (Figure S5c). Therefore, we concluded that sufficient biotinylation of target proteins by ProteasomeID can be achieved with 3 x daily biotin injections.

Using the chosen conditions (31 days of doxycycline induction and 3 x biotin injections), we enriched biotinylated proteins from all organs except the brain and analyzed them by LC-MS/MS (Figure S5d). We revealed successful enrichment of proteasome components and known interacting proteins in all the organs tested (Figure 6c and Table S5). We then compared the ProteasomeID enriched proteins identified in cultured cells and mice, and identified 116 proteins shared between HEK293T and at least one mouse organ (Figure 6d). 46 of these proteins included proteasome subunits, PIPs or members of the UPS network. In addition, we detected 70 potential novel candidate proteasome interacting proteins that were consistently identified both *in vitro* and *in vivo* (Table S5). We selected 2 of these proteins for validation by co-immunoprecipitation (Figure 6e). We expressed immunoglobulin-binding protein 1 (IGBP1) and transgelin 2 (TAGLN2) as GFP-fusions in HEK293T cells, and performed pull downs using GFP-traps. Immunoblot analysis of GFP-trap eluates confirmed co-immunoprecipitation of these proteins and endogenous PSMA4. Together these data demonstrate that known and novel proteasome interacting proteins can be identified from mouse organs using ProteasomeID.

## DISCUSSION

Our study demonstrates that proximity labeling coupled to mass spectrometry can be used to detect protein-protein interactions of the proteasome both *in vitro* and *in vivo*. Comparison to previous studies that used complementary biochemical approaches showed that ProteasomeID can retrieve a broad range of proteasome-interacting proteins in a single experiment. These include all the core proteasome subunits, activators, as well as assembly factors and components of the UPS, some of which have not been detected in previous studies. We demonstrated that ProteasomeID can be successfully implemented in mice allowing us to quantify proteasome interacting proteins in multiple mouse organs. The expression of tagged proteasomes did not negatively impact animal well-being. However, we noted that the relative levels of tagged proteasomes varied between organs, limiting the efficiency of ProteasomeID, especially in the brain. The locus used for PSMA4-miniTurbo expression (Col1a1) is known to be active in all the organs tested ^48^, however the limited penetration of food-administered doxycycline ^49^ limits the usability of our approach in the brain. In the future, crossing the ProteasomeID mouse with lines expressing the Cre recombinase under the control of specific drivers will enable the study of proteasome composition and interactions with cell type resolution *in vivo*. However, for rare cell populations, pooling organs for multiple mice might be required to achieve sufficient input material for ProteasomeID. The ProteasomeID mouse might enable future mechanistic studies on the involvement of proteasomes in malignant, inflammatory, and metabolic diseases by means of crossing with established disease models. Recently, a mouse line has been reported enabling conditional expression of a 3xFLAG tagged version of the core proteasome subunit PSMA3/α7. Similarly to our findings, tagging of the proteasome at this location did not interfere with proteasome assembly or activity, and it enabled isolation of proteasomes specifically from neurons in different models of Alzheimer’s disease ^53^.

The combination of proximity labeling and proteasome inhibition enabled the identification of known proteasome activators and endogenous substrates of the proteasome, including low abundant proteins such as transcription factors that are typically challenging to quantify by bulk proteomic analysis. Previously, approaches based on proximity labeling and mass spectrometry have been used to detect substrates of ubiquitin ligases ^54,55^ and other proteases, i.e., caspase-1 ^56^. Our work demonstrates that a similar strategy can be implemented also for proteasomes. Importantly, the measured increases of protein abundance following proteasome inhibition were considerably more pronounced in ProteasomeID than total proteome analysis. This suggests that monitoring the proteasome proximal proteome might provide higher sensitivity in the detection of proteasome substrates. Although we identified over 100 candidate substrates, the actual number of proteasome substrates is predicted to be much higher ^57^. One possible explanation for this difference is that our study only examined a single time point (4 hours) following proteasome inhibition. This short duration may have excluded substrates with slower turnover rates. To obtain a more comprehensive understanding, future studies should consider investigating multiple time points, including longer durations, to capture a broader range of substrates, including those with slower turnover rates. We envisage that ProteasomeID could be used in the future to monitor how perturbation of cellular proteostasis by different proteotoxic stress, e.g., protein aggregate formation ^25^, influences the protein degradation landscape of the proteasome, or to monitor dynamic changes in proteasome interactions and substrates, e.g., during the cell cycle, cell differentiation or cancer transformation.

Finally, we showed that ProteasomeID can directly identify known targets of PROTACs within cells. It is conceivable that the same strategy can be adapted to mouse models using the ProteasomeID mouse that we developed. This would enable assessing the efficacy of protein degraders in specific organs or cell types *in vivo.* In the future, quantification of the proteasome-associated proteome by ProteasomeID could greatly complement other existing approaches such as mass spectrometry analysis of proteolytic peptides (MAPP) ^57^ to profile the proteasome degradation landscape in physiological and disease models, and to comprehensively assess the efficacy and specificity of protein degraders.

## Acknowledgments

The authors gratefully acknowledge support from the FLI Core Facilities Proteomics, Imaging, Functional Genomics, Histology & Electron Microscopy and the Mouse Facility. The authors acknowledge Anja Baar, Christoph Kaether for support with animal experiments. AO acknowledges funding from the DFG (RTG2155 ProMoAge), the Else Kröner Fresenius Stiftung (award number: 2019_A79), the Deutsches Zentrum für Herz-Kreislaufforschung (award number: 81X2800193), the Fritz-Thyssen foundation (award number: 10.20.1.022MN), the Chan Zuckerberg Initiative Neurodegeneration Challenge Network (award numbers: 2020-221617, 2021-230967 and 2022-250618), and the NCL Stiftung. The FLI is a member of the Leibniz Association and is financially supported by the Federal Government of Germany and the State of Thuringia.

## Author contributions

Conceptualization: AB, JH, TD, JMK, AO. Investigation: AB, IH, OO, HK, ES, AM, MH, JMK. Methodology: AB, JMK. Data analysis: AB, DDF, AO. Project administration: TD, AO. Resources: AO. Supervision: JH, TD, AO. Visualization: AB, DDF, HK, AO. Writing – original draft: AB, AO. Writing – review & editing: JH, TD and JMK.

## METHODS

### Chemicals

The following chemicals were obtained from Sigma Aldrich: Dulbecco’s modified Eagle’s medium (DMEM) high glucose 4.5 g/l (D6429), L-Glutamine (G7513), biotin (B4501), PBS (D8537), complete™ Mini EDTA-free Protease Inhibitor (04693132001), Tetracycline (87128), HEPES (H3375), Sodium dodecyl sulfate (SDS) (75746), PonceauS (P7170), Sodium deoxycholate from (30970), Naphtol blue black (N3393), Adenosine triphosphate (ATP) (A2383), MG132 (M7449 and 474787), Duolink^®^ Blocking Solution and Duolink^®^ Antibody Diluent and Duolink^®^ In Situ PLA^®^ Probe Anti-Mouse MINUS (DUO92004), Duolink^®^ In Situ wash buffer A and Duolink^®^ In Situ Wash Buffer B (DUO82049), Duolink^®^ In Situ PLA® Probe Anti-Rabbit PLUS (DUO92002), Ligase from Duolink^®^ In Situ Detection Reagents Red kit and Polymerase from Duolink^®^ In Situ Detection Reagents Red kit and 5x Amplification Red buffer from Duolink^®^ In Situ Detection Reagents Red kit (DUO92008), Duolink^®^ In Situ Mounting Medium with DAPI (DUO82040), NP-40 (I8896), KCl (P3911), Iodoacetamide (I1149), Dimethyl sulfoxide (D2438), Cycloheximide (C7698).

The following chemicals were obtained from Thermo Fisher Scientific: Hygromycin B (10687010), Trypsin-EDTA (25300-062), Zeocin™ (R25001), Enhanced chemiluminescence detection kit (ECL) (32209), Sulfo-NHS-Acetate (20217), DAPI (4’,6-Diamidino-2-Phenylindole, Dihydrochloride) (D1306), Permafluor mounting medium (TA-006-FM), Poly-D-Lysine (A3890401), Fetal bovine serum (FBS) (Gibco^TM^, 10270-106) and Blasticidin (Gibco^TM^, R21001), Goat serum (Invitrogen™, 31872), Streptavidin Alexa Fluor™ 568 (Invitrogen™, S11226), EZQ^®^ Protein Quantitation Kit (Invitrogen™, R33200).

The following chemicals were obtained from Carl Roth: D(+)-biotin (3822.1), EDTA (8043.2), EGTA (3054.1), NaCl (3957.1), Triton X-100 (3051.3), Tris (4855.2), Glycerin (7533.1), β-mercaptoethanol (4227.3), Tween-20 (9127.1), Bovine serum albumin (BSA) (3737.3), Acetic acid (6755.1), Aprotinin (A162.3), Leupeptin (CN33.2), Ammonium bicarbonate (T871.2), Formic acid (4724.3), Formaldehyde (CP10.1).

The following chemicals were obtained from Biosolve: Methanol (0013684102BS), Acetone (0001037801BS), Formic acid (0006914143B5), Trifluoroacetic acid (0020234131BS), Acetonitrile (0001204102BS), 2-propanol (0016264101BS).

The following chemicals were obtained from Roche: X-tremeGENE™ 9 DNA Transfection Reagent (06365779001), Phosphatase inhibitors (04906837001), protease inhibitors (04693159001).

MgCl_2_ was obtained from Merck (8.14733.0100). Trypsin was obtained from Promega (V511 sequencing grade). LysC was obtained from Wako (125-05061 sequencing grade). Phusion^®^ High-Fidelity DNA Polymerase was obtained from NEB (New England Biolabs, M0530S). Glycine was obtained by VWR (1042011000). Turbonuclease was obtained from MoBiTec GmbH (GE-NUC10700-01). KB02-JQ1 was purchased from MedChemExpress (Cat. No. HY-129917). Urea was obtained from Bio Rad (161-0730). iRT peptides were obtained from Biognosys (iRT kit, Biognosys, Ki-3002-1).Mice food was obtained from ssniff: standard chow (ssniff, V1524-786) or with doxycycline containing food (ssniff, A153D00624). From VWR following chemicals were obtained: xylene (VWR, 28973.363), ethanol (VWR, 85830.360).

### Antibodies

Following antibodies were used: anti-FLAG^®^ M2 (1:1000, Sigma Aldrich, F3165), anti-FLAG (1:500, Sigma-Aldrich, F1804), Streptavidin HRP (1:40000, Abcam ab7403), anti-PSMA4 (1:250, NOVUS biologicals NBP2-38754), anti-SYNJ1 (1:250, Sigma Aldrich, HPA011916), anti-β-actin (1:5000, Sigma Aldrich, A5441), anti-PSMC2 (1:1000, Proteintech, 14905-1-AP), anti-c-Myc (1:1000, Abcam, ab32072), anti-Ubi-K48 (1:1000, MilliporeSigma, APU3, 05-1307), anti-ARMC6 (1:2000,Sigma Aldrich, HPA04120), anti-BRAT1 (1:10000, Abcam, ab181855), anti-GAPDH (1:200, Santa Cruz, sc-365062), anti-rabbit HRP-conjugated (1:2000, Dako, P0448), anti-mouse HRP-conjugated (1:1500, Dako P0447), anti-FLAG^®^ (1:100, Sigma Aldrich, F7425), anti-mouse-Cyanine5 (1:400, Thermo Fisher Scientific, A10524), anti-mouse Alexa Fluor 488 (1:1000, Invitrogen, A21121), Streptavidin Alexa Fluor™ 568 (1:2000, Invitrogen, S11226), anti-Proteasome 20S alpha 1+2+3+5+6+7 (1:200, Abcam, ab22674), anti-GFP (Santa Cruz, sc-9996, 1:1000), anti-BirA (1:500, Novus biologicals, NBP2-59939), anti-TIGD5 (1:1000, Proteintech, 13644-1-AP)

### Mice

Rosa26 mice (B6.Cg-Col1a1tm1(tetO-cDNA:Psma4)Mirim/J; B6.Cg-Gt(ROSA)26Sortm2 (CAG-rtTA3,-mKate2)Slowe/J) (Dow et al., 2014) were generated by Mirimus Inc. (NY, USA). All animals were housed (2-5 mice per cage) at the Leibniz Institute on Aging - Fritz Lipmann Institute, in environmentally controlled, pathogen-free animal facility with a 12 h light/ 12 h dark cycle and fed ad libitum with a standard chow or with the doxycycline containing food (625 mg/kg of dry food) for 10 or 31 days. Animals used for the procedure were 2-4 months old. Biotin (24 mg/kg body weight) or PBS were administered subcutaneously, daily for the last 3 consecutive days of the regime. At the end of the regime, mice were euthanized with CO_2_ in a CO_2_ chamber (VetTech Solutions Ltd., AN045) and the organs isolated using scissors (FST, 14090-09) and forceps (FST, 11018-12). Isolated tissues to be used for mass spectrometry analysis were washed in PBS, weighted, snap-frozen in liquid nitrogen and stored at -80°C. Isolated tissues to be used for immunofluorescence analysis were fixed in 4% formaldehyde, and embedded in paraffin for sectioning using a HistoCore Arcadia H and C (Leica). 4 µm sections were cut using a microtome HM 340E (Thermo Fisher) and placed on microscope slides (Menzel, 041300). For subsequent immunofluorescence staining, sections were rehydrated through graded alcohols using a Autostainer XL (Leica) by 2 washes for 10 min in xylene followed by 2 washes in 100% ethanol for 3 min and 1 min each, 1 min wash in 95% ethanol, 1 min wash in 70% ethanol and 50% ethanol (all v/v in water). Next, slides were washed with PBS and following this usual protocol for immunofluorescence (see below) was used.

All the procedures were conducted with a protocol approved by animal experiment license NTP-ID 00040377-1-5 (FLI-20-010) in accordance with the guidelines of the 2010/63 EU directive as well as the instructions of GV SOLAS society.

### Cell culture and treatments

FlpIn T-REx 293 cells (Thermo Fisher Scientific, R78007), referred to as HEK293T, expressing PSMA4-BirA*, PSMC2-BirA*, BirA*-PSMD3, PSMA4-miniTurbo or miniTurbo were generated as described elsewhere(Mackmull et al, 2017). Cells were grown in DMEM high glucose 4.5 g/l supplemented with 10% (v/v) heat inactivated FBS, 2 mM L-Glutamine, 15 µg/ml Blasticidin and 100 µg/ml Hygromycin B. U-2 OS cells (ATCC, HTB-96^™^; a kind gift from Pospiech lab, Leibniz Institute on Aging-Fritz Lipmann Institute) were grown in the same cell culture medium as FlpIn HEK293 T-REx cells, without antibiotics. All the cells were grown at 37°C, 5% CO_2_ and 95% humidity in a CO_2_ incubator. The parental FlpIn T-REx 293 cell line was grown in presence of 100 µg/ml Zeocin^™^ and 15 µg/ml Blasticidin. Upon generation of stable cell lines, Zeocin^™^ was replaced by 100 µg/ml Hygromycin B. For the BioID experiments, HEK293T lines were seeded at the density of approximately 1.6 x 10^4^ cells/cm^2^ and incubated for 24 h to allow cell attachment to the culture dish. The expression of BirA* or miniTurbo fusion proteins were induced by a single addition of tetracycline stock (solved in ethanol) exposing the cells to its final concentration of 1 µg/µl in total for 4 days. 2 h (miniTurbo lines) or 24 h (BirA* lines) prior to cell harvesting, 50 µM biotin was added to the culture media. For identification of proteasome substrates, PSMA4-miniTurbo or miniTurbo expressing cells were treated with 20 µM MG132 for 4 h and/or 10 µM KB02-JQ1 for 12 h. Upon treatment, cells were washed 3 x times with PBS and harvested by trypsinization (0.05% trypsin incubated for 2 min at 37°C). For each sample, a pellet corresponding to 20 million cells was collected and snap-frozen in liquid nitrogen.

### Immunoblot

Cell pellets were lysed in 50 mM HEPES pH 7.5, 5 mM EDTA, 150 mM NaCl, 1% (v/v) Triton X-100, prepared with phosphatase inhibitors and protease inhibitors for 30 min on ice. Lysates were cleared by centrifugation for 15 min at 21000 x *g* at 4°C, supernatants transferred to fresh tubes and mixed with loading buffer (1.5 M Tris pH 6.8, 20% SDS (w/v), 85% glycerin (w/v), 5% β-mercaptoethanol (v/v)). This was followed by denaturation for 5 min at 95°C. 10-20 µg of sample, determined by use of EZQ® Protein Quantitation Kit, was loaded on a 4–20% Mini-Protean^®^ TGX™ Gels (BIO-RAD) per lane and separated by SDS-PAGE. Proteins were transferred to a nitrocellulose membrane with a Trans-Blot^®^Turbo™ Transfer Starter System (BioRad, 170-4155). For high molecular weight samples (SYNJ1-BirA*, and BirA*-SYNJ1) a wet transfer method was used with Hoefer™ TE22 Mini Tank Blotting Unit (Thermo Fisher Scientific, 03-500-216), using a wet transfer buffer (25 mM Tris pH 8.3, 192 mM glycine, 15% (v/v) Methanol). Membranes were stained with PonceauS for 5 min on a shaker, washed and imaged on a Molecular Imager ChemiDoc^TM^ XRS+ Imaging system (BioRad) and destained with TBST for 1 min at room temperature (RT). After incubation for 1 h in blocking buffer (3% BSA (w/v), 25 mM Tris, 75 mM NaCl, 0.5% (v/v) Tween-20), membranes were incubated overnight at 4°C with primary antibodies diluted in blocking buffer for FLAG^®^ M2 (1:1000), Streptavidin HRP (1:40000), PSMA4 (1:250), SYNJ1 (1:250), β-actin (1:5000). This was followed by a 1 h incubation with secondary antibodies dilution matching species conjugated with HRP (1:2000, anti-rabbit; 1:1500, anti-mouse, in 0.3% BSA in TBST (w/v)). Proteins were detected using the enhanced chemiluminescence detection kit (ECL) following the manufacturer instructions. Signals were acquired on the Molecular Imager ChemiDoc^TM^XRS+ Imaging system.

For immunoblots of anti-FLAG and Streptavidin-HRP on samples from PSMA4-BirA* and PSMC2-BirA*, the cells were lysed in RIPA buffer (150 mM NaCl, 1% Triton X-100 (v/v), 0.5% sodium deoxycholate (w/v), 0.1% SDS (w/v); 50 mM Tris, pH8) prepared with phosphatase inhibitors and protease inhibitors. Samples were incubated on ice for 10 min and lysates were prepared by sonication in Bioruptor^®^ Plus sonication device (Diagenode). The following steps were performed as indicated previously, but using a different buffer to check the efficiency of the proteins to the membrane (amido black solution (0.25% (w/v), naphthol blue black, 45% (v/v) methanol, 10% (v/v) acetic acid, in milliQ water). After the ECL reaction, membranes were visualized on a CL-XPosure™ Film (Thermo Fisher Scientific, 34090), using an Amersham Hypercassette Autoradiography Cassette (RPN11648).

### Proteasome activity assay

The proteasome activity assay (PAA) was performed using the 20S proteasome activity assay kit (Millipore, APT280) following the manufacturer instructions. In short, cell pellets were thawed in ice-cold lysis buffer (50 mM HEPES pH 7.5, 5 mM EDTA, 150 mM NaCl, 1% (v/v) Triton X-100, 2 mM ATP) and left on ice for 30 min with short vortex steps every 10 min. Samples were centrifuged at 20817 x *g*, for 15 min at 4°C to remove any debris. For protein amount estimation the EZQ^®^ Protein Quantitation Kit was used. 50 μg of protein extract were incubated with fluorophore-linked peptide substrate (LLVY-7-amino-4-methylcoumarin, AMC) for 60 min at 37°C. Proteasome activity was measured by quantification of fluorescent units from cleaved AMC at 380/460 nm using a microplate reader m1000 (Tecan).

### Cycloheximide chase assay

HEK293T cells were seeded in 6 well plates (300 k cells per well). The cells were simultaneously incubated with 50 μg/ml cycloheximide for 0, 4, 8 and 12 h in the presence or absence of 20 μM MG132 proteasome inhibitor. To induce expression of the construct (PSMA4-BirA*), cells were exposed to 1 µg/µl of tetracycline. Cells were then harvested as described above. Cell pellets were then prepared by the same method and lysis buffer described in Proteasome activity assay for Western blot analysis. 10 µg of total proteins, determined by use of EZQ® Protein Quantitation Kit, per lane were separated on SDS PAGE and analyzed by immunoblotting on anti-c-myc, anti-ARMC6, anti-BRAT1, anti-TIGD5 and anti-GAPDH antibodies. The intensity of the bands were analyzed with the Image Lab™ software (v6.0.1, Bio-Rad Laboratories, Inc.).

### Native gel electrophoresis

Pellets of one million HEK293T cells containing an estimated 100 µg of total protein were collected. The pellets were then lysed in the same lysis buffer and in the same conditions as used for Proteasome activity assay. Following this, the samples were transferred into fresh tubes and 50 µg of each sample was subjected to native gel electrophoresis to reveal the various proteasome complexes (30S, 26S, 20S) using a NativePAGE™ 3 to 12% Bis-Tris gel (Invitrogen, BN1001BOX) in XCell™ SureLock™ Mini-Cell system (Invitrogen). Appropriate running buffers for native page were used (NativePAGE™ Running Buffer Kit, Invitrogen, BN2007) with addition of providing 2 mM ATP in buffers coming into direct contact with the gel. The gel was run for 60 min at 4 °C with constant 150 V, followed by voltage increase to 250V and continued to run for another 90 min. Proteins in native gels were transferred to nitrocellulose membranes for 2 h at 28V, 0.11 mA, at 4 °C, by using wet transfer buffer (48 mM Tris, 390 mM Glycine, 0.1% (w/v) SDS, 20% Methanol). The membranes were then blocked in 3% BSA, 0.5% TBST and immunoblotted with monoclonal antibodies against PSMA4 (1:250 dilution), FLAG M2 (1:1000 dilution) and PSMC2 (1:1000 dilution) overnight at 4 °C. Membranes were further incubated with horseradish peroxidase-conjugated secondary antibodies for 1 h. Proteins were detected using the Pierce™ ECL Western Blotting Substrate (Thermo Scientific, 32106) and the intensity of the bands were analyzed by the software Image Lab™ (v6.0.1, Bio-Rad).

### GFP Trap

For generation of cell lines transiently expressing either novel interactors fused to GFP or GFP control, plasmids were either bought or generated using the Gateway Technology (Invitrogen). Three 10 cm dishes of HEK293T (4 million cells/dish) were used for each candidate and prior to transfection, the medium was replaced with a transfection medium (DMEM with 2% FBS, without antibiotics). Cells were transfected with 5 µg plasmid DNA and 15 µg Polyethylenimine (PEI 25K™, Polysciences, 23966-100), both previously prepared in OptiMEM (Gibco, 11520386). The transfection medium was changed to DMEM with 10% FBS and 1% Pen-Strep 6 h post-transfection and incubated for 48 h at 3.5%, CO_2_, 37°C. Subsequently, cells were harvested by trypsinization and 20 million cell pellets were collected.

For immunoprecipitation reactions, each pellet was lysed in 200 µl of lysis buffer (20 mM Tris pH 7.5, 150 mM NaCl, 1 mM EDTA, 0.3% Triton, 10% Glycerol) containing protease and phosphatase inhibitors. Tubes were placed on ice for 30 min with vortexing every 10 min, followed by brief sonication in a Bioruptor Plus for 5 cycles (30 sec on/30 sec off) at high setting and afterwards centrifuged at 20000 x g, for 10 min at 4°C. Lysate-supernatants were transferred to a pre-cooled tube and to each of them 300 µl Dilution buffer (10 mM Tris pH 7.5, 150 mM NaCl, 0.5 mM EDTA) was added. 25 μl of GFP Trap beads (ChromoTek GFP-Trap^®^ Agarose, proteintech, gta) were equilibrated in 0.5 ml of ice-cold Dilution buffer and were spun down at 100 × g for 5-10 sec at 4°C. Beads were washed two more times with 500 μl Dilution buffer. In total, 2 mg of lysate-supernatant was added to equilibrated GFP Trap beads and were incubated for 1 h, 4°C with constant mixing. Tubes were spun at 100 × g for 5-10 sec at 4°C. GFP Trap beads were washed with 500 μl ice-cold Dilution buffer, followed by two time wash with Wash Buffer (10 mM Tris pH 7.5, 150 mM NaCl, 0.05% P40 Substitute, 0.5 mM EDTA). 80 µl of 2 x SDS-sample buffer (120 mM Tris pH 6.8, 20% glycerol, 4% SDS, 0.04% bromophenol blue) was then added to the GFP Trap beads and boiled for 5 min at 95°C. The beads were collected by centrifugation at 2500 × g for 2 min and SDS-PAGE was performed with the supernatant. Antibodies used for immunoblotting analysis: anti-PSMA4 (Novus Biologicals, NBP2-38754, 1:250), 2nd Ab: anti-rabbit (Agilent Dako, P0448, 1:2000), anti-GFP (Santa Cruz, sc-9996, 1:1000), 2nd Ab: anti-mouse (Agilent Dako, P0447, 1:1500).

### Size-exclusion chromatography (SEC)

Pellets of 80 million HEK293T cells expressing PSMA4-BirA* were collected and snap frozen in liquid nitrogen. The pellets were resuspended in 2 ml lysis buffer (50 mM HEPES pH 6.8, 1 mM MgCl_2_, 1 mM DTT, 20 mM NaCl, 5% glycerol, phosphatase inhibitors and protease inhibitors) and incubated 30 min on ice. Cell swelling and lysis was checked in 15 min intervals. Cell lysis was assisted by passage of the sample through a 27 G needle 12 times. Following this, the final concentration of NaCl was adjusted to 150 mM. The samples were then clarified by subsequent centrifugation steps as follows: (i) 500 x *g* for 5 min at 4°C, (ii) 1000 x *g* for 13 min at 4°C, and (iii) 100000 x *g* for 30 min at 4°C. The final supernatant was concentrated using 30 kDa cut-off spin filters (Merck Amicon Ultra −0.5 ml, centrifugal filters, UFC503096) to a final protein concentration of approximately 10 μg/μl measured by OD280, and further applied to size-exclusion chromatography.

SEC was performed using an ÄKTA avant (GE Äkta avant 25-1) system equipped with UV detection at 280 nm wavelength. A Yarra-SEC-4000 column (300 × 7.8 mm, pore size 500 Å, particle size 3 μm) was used with a SecurityGard™ cartridge GFC4000 4 × 3.0 mm ID as a guard column. Running conditions were 4°C, a flow rate of 0.5 ml/min and run time of 40 min. The mobile phase contained 50 mM HEPES, pH 6.8, 1 mM MgCl_2_, 1 mM DTT, 150 mM NaCl, and 5 mM ATP. A control sample (Phenomenex, ALO-3042) was injected prior to each sample to verify column performance. 100 μl samples from 10 mg/ml lysate solution were injected, corresponding to 1 mg protein extract on column. Fractions (200 μl each) were collected along with the LC (liquid chromatography) separation directly in the SDS buffer, to a final concentration of 4%. Thirty-six fractions were further processed for LC-MS/MS analysis. Of these 36 fractions the first and last two fractions were pooled.

### Preparation of SEC fractions for mass spectrometry analysis

The SEC fractions were further processed by addition of DTT (50 mM) in 100 mM HEPES at pH 8, boiled for 5 min at 95°C, followed by sonication (Diagenode Bioruptor^®^ Plus) for 10 cycles (30 sec on/60 sec off) at 4°C. The samples were then centrifuged at 3000 x *g* for 5 min at RT, and the supernatant transferred to 2 ml tube. This was followed by alkylation with 20 mM iodoacetamide (IAA) for 30 min at RT in the dark. Protein amounts were confirmed by SDS–PAGE (4%). Protein samples in the collected fractions ranged from 10 - 100 µg. Proteins were precipitated overnight at −20°C after addition of a 4 × volume of ice-cold acetone. Thereafter, the samples were centrifuged at 20800 x *g* for 30 min at 4°C and the supernatant carefully removed. Pellets were washed twice with 1 ml ice-cold 80% (v/v) acetone and then centrifuged with 20800 x *g* at 4°C. The samples were air-dried before addition of 120 μl digestion buffer (3 M urea, 100 mM HEPES, pH8). Samples were resuspended by sonication (as above) and LysC was added at 1:100 (w/w) enzyme:protein ratio. The samples were then digested for 4 h at 37°C (1000 x rpm for 1 h, then 650 x rpm, Eppendorf ThermoMixer^®^C). Samples were then diluted 1:1 with milliQ water, and trypsin added at the same enzyme to protein ratio and further digested overnight at 37°C (650 x rpm). Consequently, digests were acidified by the addition of TFA to a final concentration of 2% (v/v) and then desalted with a Waters™ Oasis^®^ plate (HLB μElution Plate 30 μm, Waters, 186001828BA) with slow vacuum. Therefore, the columns were conditioned three times with 100 μl solvent B (80% (v/v) acetonitrile, 0.05% (v/v) formic acid) and equilibrated three times with 100 μl solvent A (0.05% (v/v) formic acid in Milli-Q water). The samples were loaded, washed 3 times with 100 μl solvent A, and then eluted with 50 μl solvent B. The eluates were dried in a vacuum concentrator.

### BioID affinity purification

Cell pellets were thawed on ice and resuspended in 4.75 ml BioID lysis buffer (50 mM Tris pH 7.5, 150 mM NaCl, 1 mM EDTA, 1 mM EGTA, 1% (v/v) Triton X-100, 1 mg/ml aprotinin, 0.5 mg/ml leupeptin, 250 U turbonuclease, 0.1% (w/v) SDS), followed by 1 h incubation in the rotator mixer (STARLAB RM Multi-1) (15 rpm) at 4°C to aid the lysis. Samples were then briefly sonicated in a Bioruptor Plus for 5 cycles (30 sec on/30 sec off) at high setting and afterwards centrifuged at 20817 x *g*, for 30 min at 4°C to remove any debris.

Mouse organs were thawed and transferred into Precellys^®^ lysing kit tubes (Bertin Instruments, 431-0170, Keramik-kit 1.4/2.8 mm, 2 ml (CKM)) containing 1 ml of PBS supplemented with 1 tab of complete™, EDTA-free Protease Inhibitor per 50 ml. For homogenization, organs were shaken twice at 6000 x rpm for 30 sec, centrifuged at 946 x *g* at 4°C for 5 min, and the resulting homogenate was transferred to a new tube. Based on the estimated protein content (5% of fresh tissue weight for liver, 8% for heart and kidney and 20% for muscle), homogenates corresponding to 4 mg protein were processed for further BioID affinity purification. This entailed cell lysis of the homogenates by means of BioID lysis buffer.

Streptavidin coated Sepharose beads (Merck, GE17-5113-01) were acetylated by two successive treatments with 10 mM Sulfo-NHS-Acetate for 30 min at RT. The reaction was then quenched with 1 M Tris pH 7.5 (1:10 v/v) and the beads were washed three times with 1 x PBS and centrifuged at 2000 x *g* for 1 min at RT. Cleared lysates were transferred to new tubes, 50 µl of acetylated beads added, and samples were incubated for 3 h on the rotator (15 rpm) at 4°C. This was followed by centrifugation at 2000 x *g* for 5 min at 4°C and removal of 4.5 ml of the supernatant from each sample. Remaining sample with the beads at the bottom was transferred to a Pierce Spin Column Snap Cap column (Thermo Fisher Scientific, 69725) and the tubes were additionally rinsed with a lysis buffer and added to the spin column. Beads were then washed on the column with a lysis buffer, followed by 3 washes with freshly prepared 50 mM ammonium bicarbonate (AmBic), with the pH adjusted to 8.3. The bottom of the columns were closed with a plug and beads transferred to fresh 2 ml tubes by means of 3 x 300 µl 50 mM AmBic, pH 8.3. The samples were then centrifuged at 2000 x *g* for 5 min at 4°C and the content of each tube was removed, leaving 200 µl in the tube. 1 µg of LysC was added and incubated at 37°C for 16 h shaking at 500 x rpm. The samples were then centrifuged at 2000 x *g* for 5 min at room temperature and the content of the tubes were transferred to Pierce Spin Column Snap Cap columns. The digested peptides were eluted with two times 150 μl of freshly made 50 mM AmBic. To elute biotinylated peptides still bound to the beads, 150 μl of 80% ACN and 20% TFA was added, briefly mixed, and rapidly eluted. This elution step was repeated twice and the eluates merged. Following elution, 0.5 μg of trypsin was added to the AmBic elutions and digestion continued for an additional 3 h with mixing at 500 x rpm and 37°C. Digested AmBic elutions were then dried down in a vacuum concentrator, resuspended in 200 µl 0.05% (v/v) formic acid in milliQ water and sonicated in a Bioruptor Plus (5 cycles with 1 min on and 30 sec off with high intensity at 20°C). ACN/TFA elutions were dried down in a vacuum concentrator until approximately 50 µl were left, and 50 µl of 200 mM HEPES pH 8.0 were added to the samples and pH adjusted to 7-9. 0.5 μg of trypsin were then added and digestion continued for an additional 3 h with mixing at 500 x rpm at 37°C. Digested peptides were acidified with 10% (v/v) trifluoroacetic to pH <3. Both AmBic and ACN/TFA elutions were desalted using Macro Spin Column C18 columns (Thermo Scientific™, Pierce™, 89873) following manufacturer’s instructions and dried down in a vacuum concentrator.

### LC-MS/MS data acquisition

Prior to analysis, samples were reconstituted in mass spectrometry (MS) Buffer (5% acetonitrile, 95% Milli-Q water, with 0.1% formic acid) and spiked with iRT peptides. Peptides were separated in trap/elute mode using the nanoAcquity MClass Ultra-High Performance Liquid Chromatography system (UPLC) or nanoAcquity UPLC system (Waters) equipped with a trapping (nanoAcquity Symmetry C18, 5 μm, 180 μm × 20 mm) and an analytical column (nanoAcquity BEH C18, 1.7 μm, 75 μm × 250 mm). Solvent A was water with 0.1% formic acid, and solvent B was acetonitrile with 0.1% formic acid. 1 µl of the sample (∼1 μg on column) was loaded with a constant flow of solvent A at 5 μl/min onto the trapping column. Trapping time was 6 min. Peptides were eluted via the analytical column with a constant flow of 0.3 μl/min. During the elution, the percentage of solvent B increased in a nonlinear fashion from 0–40% in 90 min (120 min for total proteome of mouse organs). Total run time was 115 min (145 min) including equilibration and conditioning. The LC was coupled to an Orbitrap Fusion Lumos (Thermo Fisher Scientific) using the Proxeon nanospray source or to an Orbitrap Q-Exactive HFX (Thermo Fisher Scientific) for BioID experiments from HEK293T cells, or to an Orbitrap Exploris 480 (Thermo Fisher Scientific) for BioID experiments combined with PROTAC treatment. The peptides were introduced into the mass spectrometer via a Pico-Tip Emitter 360-μm outer diameter × 20-μm inner diameter, 10-μm tip (New Objective) heated at 300°C, and a spray voltage of 2.2 kV was applied. For data acquisition and processing of the raw data Tune version 2.1 and Xcalibur 4.1 (Orbitrap Fusion Lumos), Tune 2.9 and Xcalibur 4.0 (Orbitrap Q-Exactive HFX) and Tune 3.1 and Xcalibur 4.4 (Orbitrap Exploris 480) were employed.

#### DDA (Data-dependent acquisition)

SEC fractions, mouse BioID as well as BioID of PSMA4 and PSMC2 were analyzed using DpD (DDA plus DIA). Here, data from a subset of conditions were first acquired in DDA mode to contribute to a sample specific spectral library. Full scan MS spectra with mass range 375-1500 m/z (using quadrupole isolation) were acquired in profile mode in the Orbitrap with resolution of 60,000 FWHM. The filling time was set at a maximum of 50 ms with a limitation of 2 x 10^5^ ions. The “Top Speed” method was employed to take the maximum number of precursor ions (with an intensity threshold of 5 x 10^5^) from the full scan MS for fragmentation (using HCD collision energy, 30%) and quadrupole isolation (1.4 Da window) and measurement in the Orbitrap, with a cycle time of 3 seconds. The MIPS (monoisotopic precursor selection) peptide algorithm was employed. MS/MS data were acquired in centroid mode in the Orbitrap, with a resolution of 15,000 FWHM and a fixed first mass of 120 m/z. The filling time was set at a maximum of 22 ms with a limitation of 1 x 10^5^ ions. Only multiply charged (2+ - 7+) precursor ions were selected for MS/MS. Dynamic exclusion was employed with maximum retention period of 15 sec and relative mass window of 10 ppm. Isotopes were excluded.

#### DIA (Data-independent acquisition)

The DIA data acquisition was the same for both directDIA and DpD. Full scan mass spectrometry spectra with mass range 350–1650 m/z were acquired in profile mode in the Orbitrap with resolution of 120,000 FWHM. The default charge state was set to 3+. The filling time was set at a maximum of 60 ms with a limitation of 3 × 10^6^ ions. DIA scans were acquired with 34 mass window segments of differing widths across the MS1 mass range. Higher collisional dissociation fragmentation (stepped normalized collision energy: 25, 27.5, and 30%) was applied and MS/MS spectra were acquired with a resolution of 30,000 FWHM with a fixed first mass of 200 m/z after accumulation of 3 × 10^6^ ions or after filling time of 35 ms (whichever occurred first). Data was acquired in profile mode.

### LC-MS/MS data analysis

DpD (DDA plus DIA) libraries were created by searching both the DDA runs and the DIA runs using Spectronaut Pulsar (v 13-15, Biognosys). The data were searched against species specific protein databases (*Homo sapiens*, reviewed entry only (16,747 entries), release 2016_01 or *Mus musculus*, entry only (20,186), release 2016_01 respectively) with a list of common contaminants appended. The data were searched with the following modifications: carbamidomethyl (C) as fixed modification, and oxidation (M), acetyl (protein N-term), and biotin (K) as variable modifications. A maximum of 2 missed cleavages was allowed. The library search was set to 1% false discovery rate (FDR) at both protein and peptide levels. This library contained 79,732 precursors, corresponding to 4,730 protein groups for SEC fractions, 77,401 precursor, corresponding to 5,125 protein groups for BioID on PSMA4 and PSMC2 using Spectronaut protein inference. All other BioID experiments were processed using the directDIA pipeline in Spectronaut Professional (v.13-17). The data were searched against a species specific (*Mus musculus* and *Homo sapiens*, as described above) with a list of common contaminants appended. BGS factory settings were used with the exception of: variable modifications = acetyl (protein N-term), biotin (K), oxidation (M). SEC-MS experiments were processed using Spectronaut v.13 with default settings except: Proteotypicity Filter = Only Protein Group Specific; Major Group Quantity = Median peptide quantity; Major Group Top N = OFF; Minor Group Quantity = Median precursor quantity; Minor Group Top N = OFF; Data Filtering = Qvalue sparse; Imputing Strategy = No imputing; Cross run normalization = OFF.

PSMA4 and PSMC2 BioID experiments were processed using Spectronaut v.13 with default settings except: Proteotypicity Filter = Only Protein Group Specific; Major Group Quantity = Median peptide quantity; Major Group Top N = OFF; Minor Group Quantity = Median precursor quantity; Minor Group Top N = OFF; Data Filtering = Qvalue percentile (0.5); Imputing Strategy = No imputing; Normalization Strategy = Global Normalization; Normalize on = Median; Row Selection = Qvalue sparse.

Mouse BioID, PSMD3 BioID and BioID experiments combined with PROTAC treatment were processed using Spectronaut v.15, v.17 and v18 with default settings except: Proteotypicity Filter = Only Protein Group Specific; Major Group Quantity = Median peptide quantity; Major Group Top N = OFF; Minor Group Quantity = Median precursor quantity; Minor Group Top N = OFF; Data Filtering = Qvalue percentile (0.2); Imputing Strategy = Global imputing; Normalization Strategy = Global Normalization; Normalize on = Median; Row Selection = Qvalue complete.

For all the BioID experiments, differential abundance testing was performed in Spectronaut using a paired t-test between replicates. P values were corrected for multiple testing multiple testing correction with the method described by Storey ^58^. The candidates and protein report tables were exported from Spectronaut and used for further data analysis using R and RStudio server.

### Logistic regression classifier for detecting ProteasomeID enriched proteins

To identify ProteasomeID enriched proteins, we trained a logistic regression binary classifier. The classifier was trained using known proteasome members as positive class, and mitochondrial matrix proteins as negative. Mitochondrial matrix proteins are not expected to interact directly with the proteasome under homeostatic conditions. To distinguish between these two classes, we performed prediction using an enrichment score derived from multiplying the average log2 ratio and the negative logarithm of the q-value obtained from a differential protein abundance analysis performed against the BirA* control line. Before analysis, any missing data points were removed from the dataset. To assess the performance of the binary classifier, and optimize its parameters, a 10-fold cross-validation approach was adopted. The dataset was randomly partitioned into ten subsets, with nine subsets used for training and one subset for validation in each iteration. This process was repeated thirty times, and results were averaged using the mean value to ensure the robustness of the results. Logistic regression was employed as the classification method using the caret package in R ^59^. To determine an optimal threshold for classification, the false positive rate (FPR) was set at 0.05. The threshold yielding an FPR closest to the target value of 0.05 was selected as the final classification threshold. Following model training and threshold selection, the classifier was applied to predict the class labels of additional proteins not used in the training process. The enrichment score and class labels for the new data were provided as input to the trained model. All statistical analyses were performed using R version 4.1.3. The pROC ^60^ and caret ^59^ packages were employed for ROC analysis and logistic regression, respectively.

### Immunofluorescence

Cells were grown on autoclaved coverslips (Carl Roth, YX03.1), coated with Poly-D-Lysine. Coverslips were placed individually in 12-well plates (Lab solute, 7696791), and 25 k cells were seeded per well. Cells were washed three times with 1 x PBS, fixed in 4% formaldehyde (v/v) in PBS for 10 min at RT, washed 3 x 5 min with 1 x PBS and permeabilized with permeabilization buffer (0.7% Triton X-100, in 1 x PBS) at room temperature for 15 min. Washing with PBS was repeated 2 x 5 min and samples were incubated with blocking solution (10% (w/v) BSA, 10% (v/v) Triton X-100, 5% (v/v) goat serum) for 10 min at RT. The coverslips were incubated with primary antibody anti-FLAG®M2 (1:100, Sigma Aldrich, mouse F3165) or anti-FLAG (1:500, Sigma-Aldrich, mouse, F1804) at 4°C overnight. After washing 3 x 5 min with PBS/PBST (first with PBS, second with PBS + 0.2% (v/v) Tween 20, third with PBS) the secondary fluorescence-labeled antibody (goat anti-mouse IgG (H+L) - Cyanine5, 1:400 in blocking solution or goat anti-mouse IgG g1 Alexa Fluor 488, 1:1000 in blocking solution) and fluorescently labeled streptavidin, 1:2000 in blocking solution) were incubated for 30 min at 37°C. After 3 x 5 min with PBS/PBST (first with PBS, second with PBS + 0.2% (v/v) Tween 20, third with PBS), nuclei were stained with DAPI (4’,6-Diamidino-2-Phenylindole, Dihydrochloride, 0.02 μg/μl in PBS) at RT for 10 min and washed again with PBS 2 x 5 min. Frozen sections were mounted in Permafluor mounting medium using glass slides (041300, Menzel) and dried at room temperature overnight. All samples were stored at 4°C in the dark until further analysis by microscopy. Immunofluorescence microscopy was performed with an Axio Imager (Z2 using a Plan-Apochromat 63 x / 0.8 M27 Objective) and analyzed with the software Zen 2 Blue Edition (Carl Zeiss Microscopy GmbH).

### Immunohistochemistry

5 μm sections were deparaffinized by xylene twice 5 min each and rehydrated by 100%, 90%, 70% ethanol for 5 min each and washed with tap water for 10 min. Sections were treated with antigen unmasking solution sodium citrate buffer (Vector, H-3300) in the microwave at 800W (3 min) followed by 400W (10 min) for antigen retrieval. Sections were cooled at room temperature and washed three times with PBS, 5 minutes each. Sections were blocked for the endogenous peroxidase activity by 0.03% H_2_O_2_ in methanol for 30 minutes at RT. Sections were rinsed with PBS three times and blocked with 5% BSA for 1 h at RT. Excess serum was tipped off and sections were incubated with primary antibody (anti-FLAG, Sigma F7425, 1:100 diluted in 1% BSA), overnight in a humid chamber at 4°C. The next day sections were brought to room temperature and washed three times with PBS, 5 min each. Sections were incubated with biotinylated rabbit antibody diluted in 1% BSA/PBS +0.01% Tween20 for 1 h. Sections were washed three times with PBS, 5 min each and incubated with a mixture from VECTASTAIN® Elite® ABC HRP Kit (PK-6100) for 30 min. It was prepared at least 20 min prior to use, one drop of solution A and one drop of solution B were mixed into 2.5 ml of ABC dilution buffer, vortexed and used. Sections were rinsed three times with PBS, 5 min each and developed by ImmPACT® NovaRED® Peroxidase (HRP) Substrate (SK-4800) according to manufacturer’s instructions until the reddish brown color visibly appeared while examining the sections under the microscope. The reaction was stopped by immersing the sections in the water for 5 min. Sections were counterstained with hematoxylin for 30 sec, rinsed in tap water, and dehydrated with 70%, 90% and 100% ethanol 30 sec each and with xylene for 1 min. Slides were mounted with xylene based mounting medium, air dried and stored in a cold and dry place for further analysis. Images were taken with an Axio Imager M2 microscope (Zeiss) equipped with an AxioCam MRc5 (Zeiss), with a 20 x objective.

### Molecular visualization and structure analysis

For visualization of proteasome complexes UCSF Chimera program (version 1.13.1) was used. The three-dimensional structural data of macromolecular complexes of proteasome were downloaded from the Protein Data Bank (PDB) database (5T0C). For the analysis of the enrichment of proteasome subunits in BioID protocol, data sets with fold change information were used and filtered the following way: q value < 0.05, number of identified unique peptides per protein ≥ 2. The intensity of the proteasome subunit coloring used was directly dependent on the fold change of the identified subunit in the BioID affinity purification.

## Data availability

Mass spectrometry proteomics data have been deposited to the ProteomeXchange Consortium via the PRIDE ^61^ partner repository and they are accessible with the identifier:

PXD032833 following login credentials Username: password: UeTUvffr, for the HEK293T ProteasomeID dataset for PSMA4 and PSMC2;

PXD034874 following login credentials Username: password: JSKJIvmX, for the HEK293T size exclusion chromatography dataset;

PXD033008 following login credentials Username: password: eRoFg8LW, for the MG132 ProteasomeID dataset;

PXD032976 following login credentials Username: password: kcQb7byl for the PROTAC ProteasomeID dataset;

PXD034965 following login credentials Username: password:

DHtFE3zX, for the MG132 whole cell proteome dataset;

Mass spectrometry proteomics data have been deposited to ProteomeXchange Consortium via the MassIVE partner repository and they are accessible with the identifier:

MSV000092396 following login credential Username: MSV000092396_reviewer password: a, for the mouse ProteasomeID dataset

MSV000092407 following login credential Username: MSV000092407_reviewer password: proteasomeID, for the HEK293T ProteasomeID for PSMD3

## SUPPLEMENTARY INFORMATION

**Figure S1:**
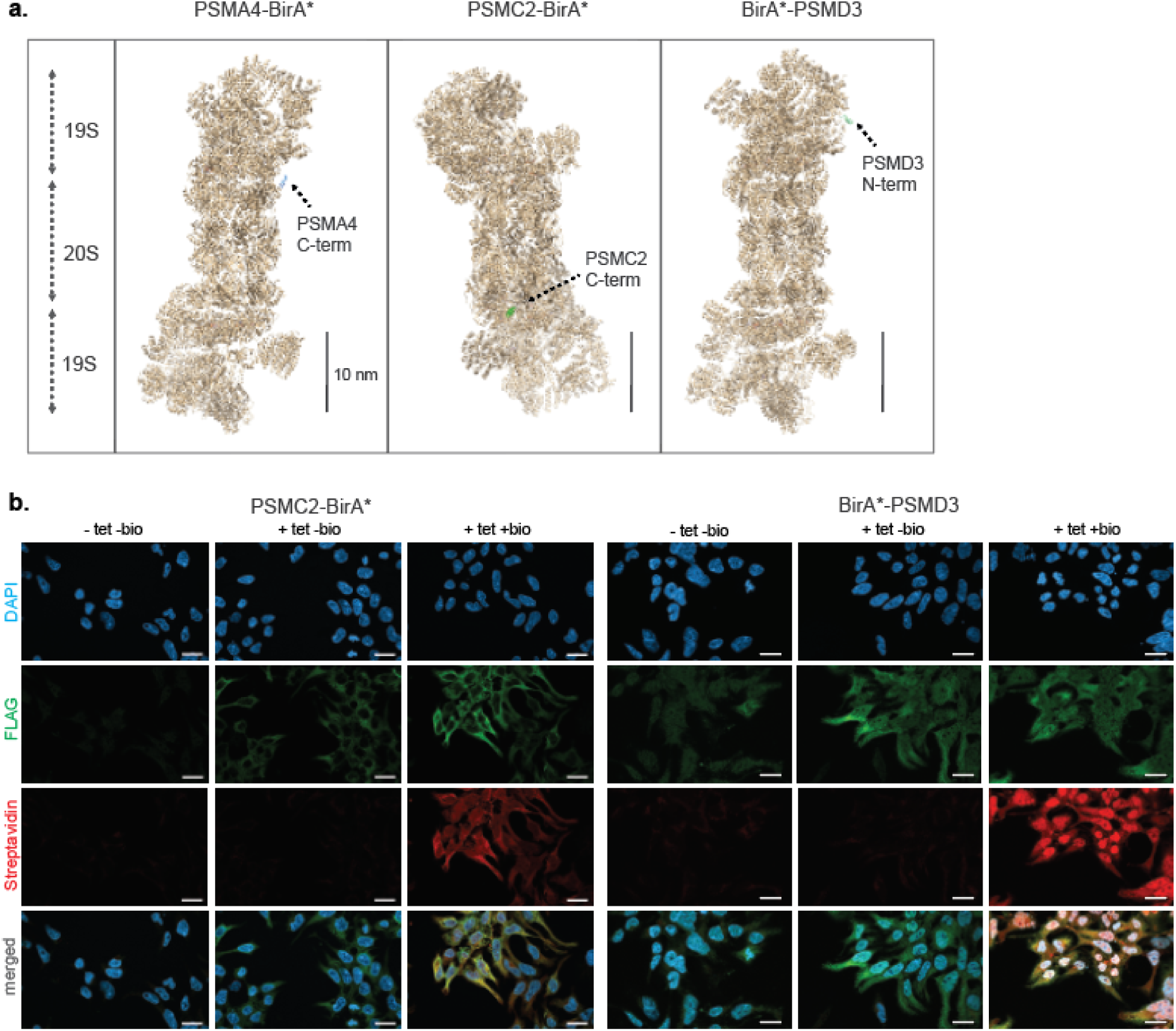
Further characterization of PSMC2-BirA*,BirA*-PSMD3 expressing cell lines. a. Schematic representation of proteasome structure with the biotin ligase-fused to N- or C-termini of the respective bait proteins highlighted in color. Scale bar = 10 nm. The proteasome structure depicted was obtained from the PDB:5T0C model of the human 26S proteasome ^62^ and rendered using Chimera ^63^. b. Immunofluorescence analysis of PSMC2-BirA*-FLAG and BirA*-FLAG-PSMD3 HEK293T cell lines 4 days after seeding without addition of any substance (-tet -bio), with addition of only tetracycline for 4 days (+tet -bio) or with addition of both tetracycline for 4 days and biotin for 1 day (+tet +bio). Scale bar = 20 µm.

**Figure S2:**
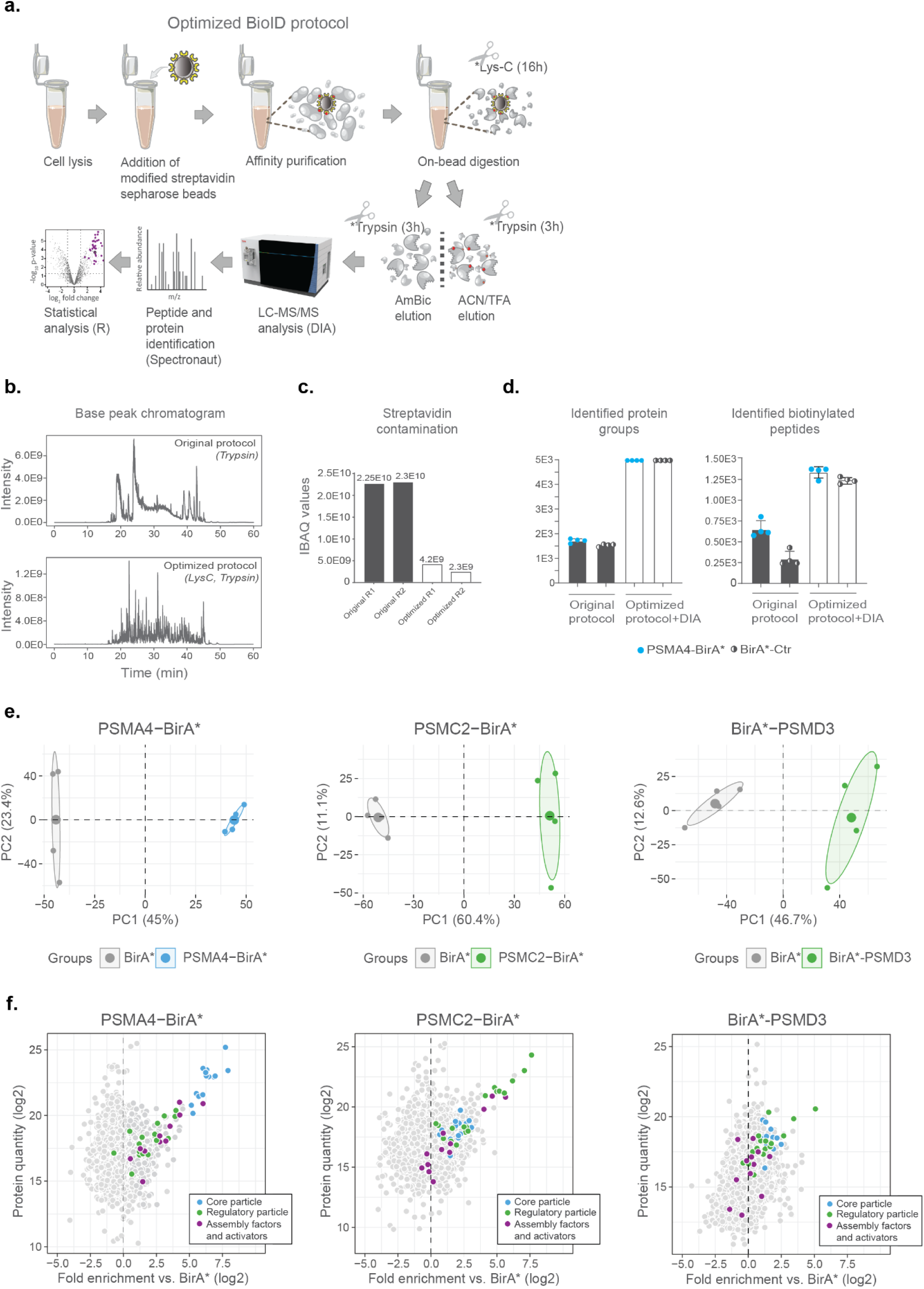
Optimization of BioID workflow and ProteasomeID cell lines evaluation. a. Scheme of optimized BioID protocol. The original protocol by Mackmull et al. ^36^ was optimized by (i) acetylation of lysines on streptavidin prior to pull-down; (ii) replacement of trypsin by LysC for on-bead digestion. Following on-bead digestion, two sequential elutions with ammonium bicarbonate (AmBic) and acetonitrile (ACN) / trifluoroacetic acid (TFA) are performed and eluates are further digested off-beads with trypsin. (iii) Peptides from AmBic and ACN/TFA eluates are then analyzed by Data Independent Acquisition (DIA) mass spectrometry. b. Representative base peak chromatograms of AmBic elutions obtained from the original (upper panel,) and modified (lower panel) BioID protocol. The replacement of trypsin with LysC together with the acetylation of streptavidin drastically reduces the contamination by streptavidin-derived peptides. c. Quantification of streptavidin in AmBic elutions obtained from the original (T: trypsin) and modified (LT: LysC followed by trypsin) BioID protocol. Streptavidin quantification was based on iBAQ values ^65^ obtained label free mass spectrometry analysis. Two representative replicates (R1, R2) are shown for each condition. d. Bar plots of the number of identified protein groups and biotinylated peptides obtained using the original vs. optimized BioID protocols. Data were obtained from cell lines expressing PSMA4-BirA* (light blue dots) or BirA* (black/white dots). The number of identified protein groups was obtained from AmBic samples, while the number of biotinylated peptides was derived from ACN/TFA samples. n = 4 biological replicates, error bars indicate standard deviation of the mean. e. Principal component analysis (PCA) of ProteasomeID cell lines samples based on the abundance of all proteins identified by label-free mass spectrometry. The smaller dots represent individual samples and the larger dots the centroids of each age-matched group. Ellipses represent 95% confidence intervals. The percentage of variance explained by the first two PC axes is reported in the axis titles. f. MA plots of proteins enriched by streptavidin pull-down and analyzed by DIA mass spectrometry from different cell lines. Highlighted in color are proteasome members or assembly factors and activators. Data were obtained from n = 4 biological replicates.

**Figure S3:**
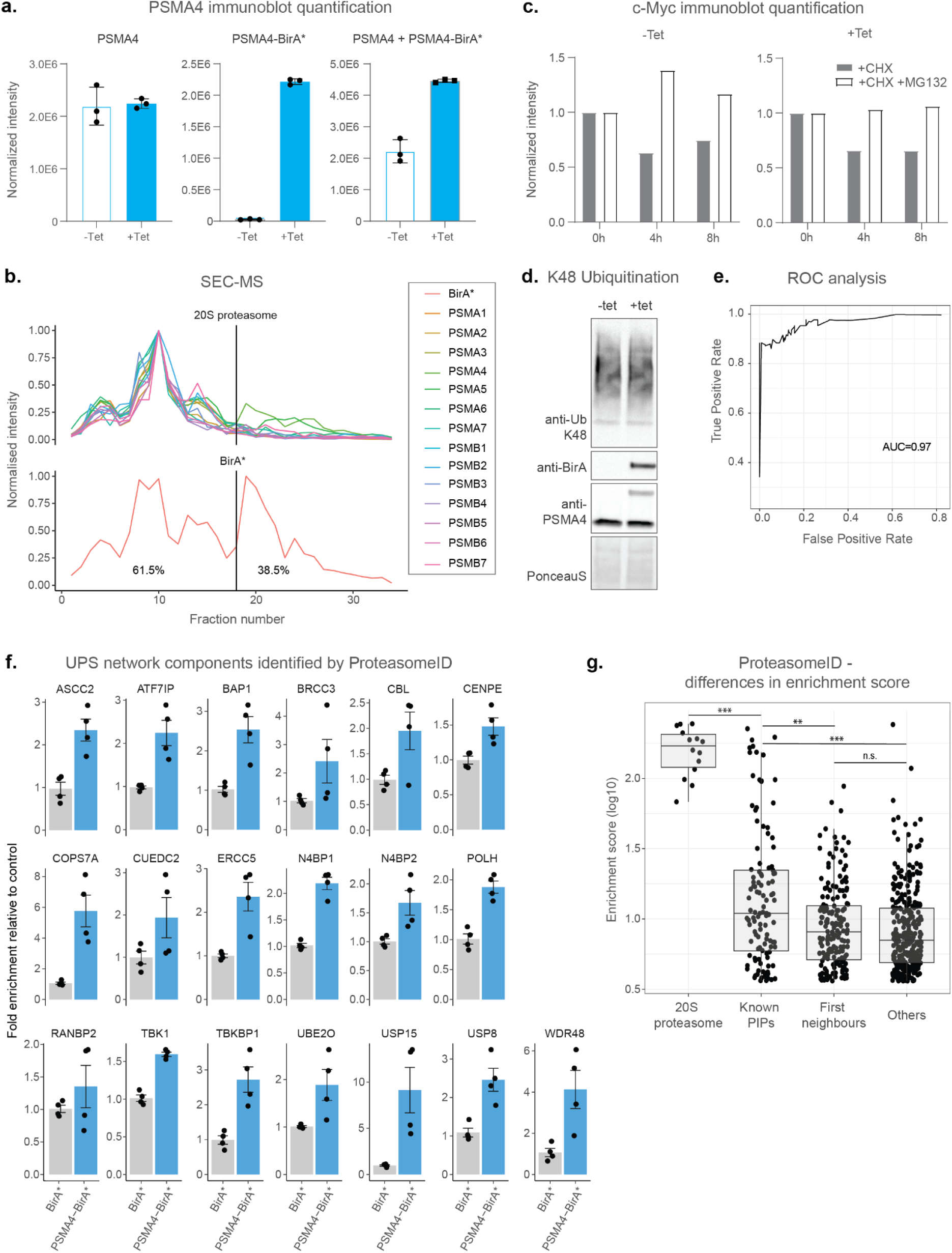
ProteasomeID cell line validation. a. Densitometric quantification of immunoblot depicted in figure 2a. Ponceau staining was used as loading control for normalization. n = 3 biological replicates, error bars indicate standard deviation of the mean. b. Size exclusion chromatography (SEC) analysis of lysates from HEK293T cells stably expressing PSMA4-BirA* following 24 h incubation with tetracycline. SEC fractions were analyzed by DIA mass spectrometry and elution profiles were built for each protein using protein quantity values normalized to the sum of quantities across all fractions. Depicted in the upper panel are the elution profiles of 20S proteasome components. The lower panel depicts the elution profile of BirA*. The perpendicular dashed line depicts a cut off for fully assembled proteasome complexes. c. Densitometric quantification of immunoblot depicted in Figure 2f. Ponceau staining was used as loading control followed by band intensity normalization to zero hour, untreated samples (which were set to 1). Tet = tetracycline, CHX = cycloheximide. d. Immunoblot comparing the levels of K48 ubiquitylated proteins from PSMA4-BirA* cells treated with (+tet) or without (-tet) tetracycline. Afterwards, expression of PSMA4-BirA* was verified by blotting against BirA* and PSMA4. Ponceau staining was used as loading control. e. ROC curve of the classifier used to define ProteasomeID enriched proteins. f. Bar plots comparing the levels of enrichment obtained in ProteasomeID experiment for members of the UPS network not identified in previous interaction studies. Enrichment levels were normalized to the levels detected in BirA* control cell line which was set to 1. Protein quantities were derived from DIA mass spectrometry data. Data are shown as mean ± standard error from n = 4 biological replicates. g. Differences in enrichment score (compared to BirA* control) between proteins of the 20S proteasome, known PIPs, first neighbors of known PIPs and other enriched proteins. The enrichment scores were obtained from DIA mass spectrometry data from n = 4 biological replicates. P-values were derived from Wilcoxon rank sum test: *** p < 0.001, ** p < 0.01, n.s. p > 0.05

**Figure S4:**
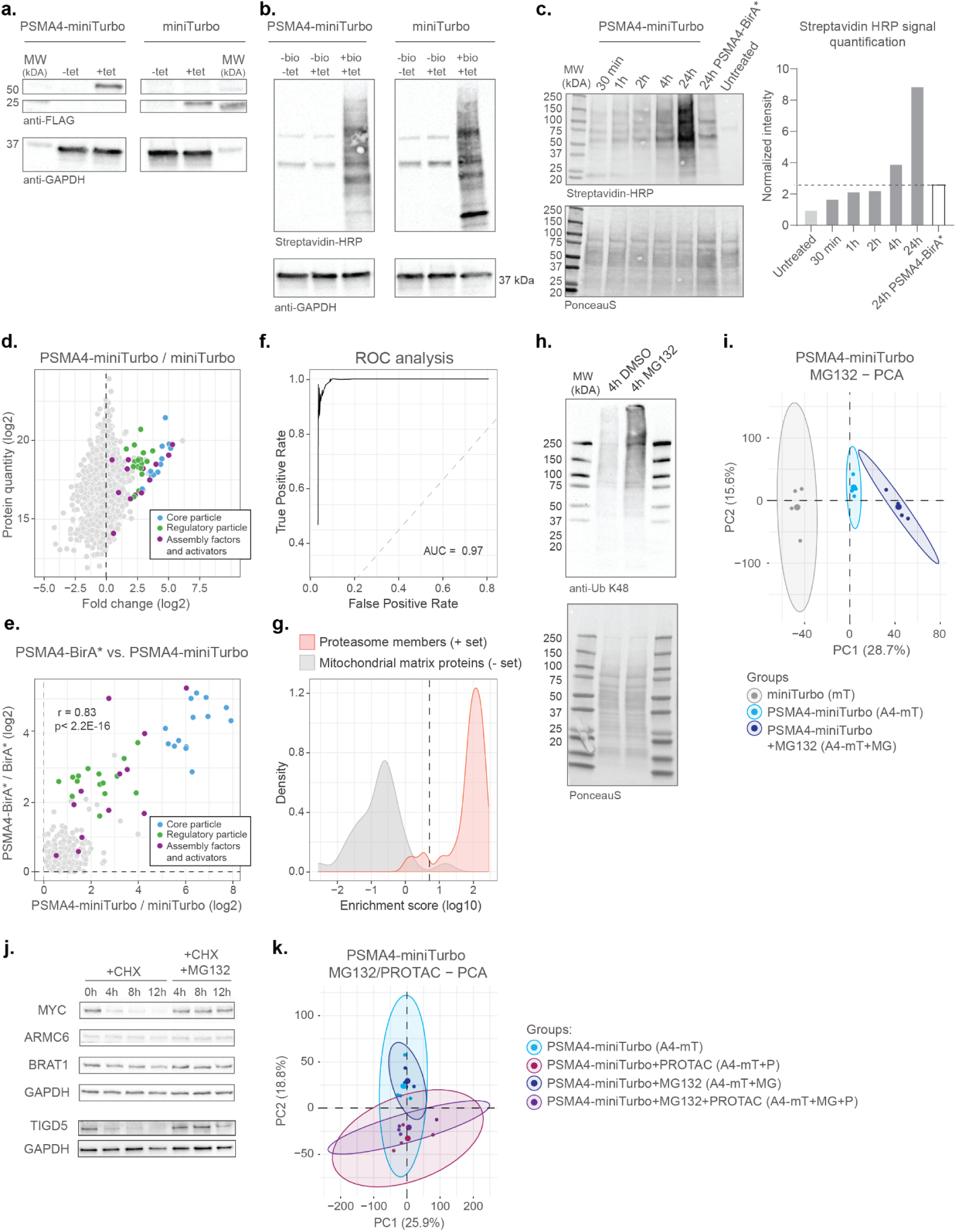
Validation of PSMA4-miniTurbo cell line and application of ProteasomeID for detecting endogenous and PROTAC-induced proteasome substrates. a. Immunoblot of miniTurbo fusion proteins performed on lysates collected from HEK293T cells stably transfected with PSMA4-miniTurbo-FLAG or miniTurbo-FLAG following 4 days of incubation with (+tet) or without (−tet) tetracycline. Immunoblot against GAPDH was used as loading control. b. Streptavidin-HRP immunoblot following induction of miniTurbo fusion proteins with tetracycline and supplementation of biotin for 2 h. Immunoblot against GAPDH was used as loading control. c. Streptavidin-HRP immunoblot following induction of miniTurbo fusion proteins with tetracycline and supplementation of biotin for indicated times. For the sample used as negative control biotin supplementation was omitted (-biotin). Ponceau staining was used as loading control. Bar plots on the left depict densitometric quantification of the immunoblot. Samples were normalized to the band intensity of PSMA4-BirA* sample not supplemented with biotin (untreated sample). d. MA plot of proteins enriched by streptavidin pull-down and analyzed by DIA mass spectrometry from PSMA4-miniTurbo and miniTurbo control cell lines. Data were obtained from n = 4 biological replicates. e. Comparison of log2 fold changes for streptavidin-enriched proteins from PSMA4-BirA* and PSMA4-miniTurbo compared to their respective controls. Proteins significant (Q value < 0.05) and displaying a log2 fold change > 0 in both comparisons were considered for the analysis. f. ROC analysis of the classifier used to define ProteasomeID (PSMA4-miniTurbo) enriched proteins. g. Distribution of enrichment scores for PSMA4-miniTurbo enriched proteins. Calculated by the classifier algorithm for proteasome subunits (set of true positives) and mitochondrial matrix proteins (set of true negatives). The dashed vertical line indicates the enrichment score cut-off to define ProteasomeID enriched proteins at FPR < 0.05. h. Immunoblot for K48 ubiquitylated proteins from PSMA4-miniTurbo cells treated with 20 µM MG132 for 4h. As a negative control the same cell line was treated in the same way with DMSO only. Ponceau staining was used as loading control. i. Principal component analysis (PCA) of ProteasomeID data obtained from cell lines expressing PSMA4-miniTurbo and control (miniTurbo), and PSMA4-miniTurbo following exposure to proteasome inhibitor MG132. The smaller dots represent individual samples and the larger dots the centroids of each group. Ellipses represent 95% confidence intervals. The percentage of variance explained by the first two principal components (PC) axes is reported in the axis titles. n = 4, biological replicates. j. Cycloheximide-chase experiment on stability of 3 potential novel proteasome substrate proteins. PSMA4-BirA*cells were incubated with 50 μg/ml cycloheximide (CHX) for the indicated times in the presence or absence of MG132 (20 μM) and tetracycline (1 µg/µl). Cell lysates were then prepared for Western blot analysis of steady-state levels of c-Myc, ARMC6, and BRAT1 and TIGD5. c-Myc was used as a positive control as it is a well known proteasome substrate. Tet = tetracycline, CHX = cycloheximide. k. Principal component analysis (PCA) of ProteasomeID data obtained from cells expressing PSMA4-miniTurbo exposed to the proteasome inhibitor MG132 and/or the PROTAC KB02-JQ1. The smaller dots represent individual samples and the larger dots the centroids of each group. Ellipses represent 95% confidence intervals. The percentage of variance explained by the first two principal components (PC) axes is reported in the axis titles. n = 4, biological replicates.

**Figure S5:**
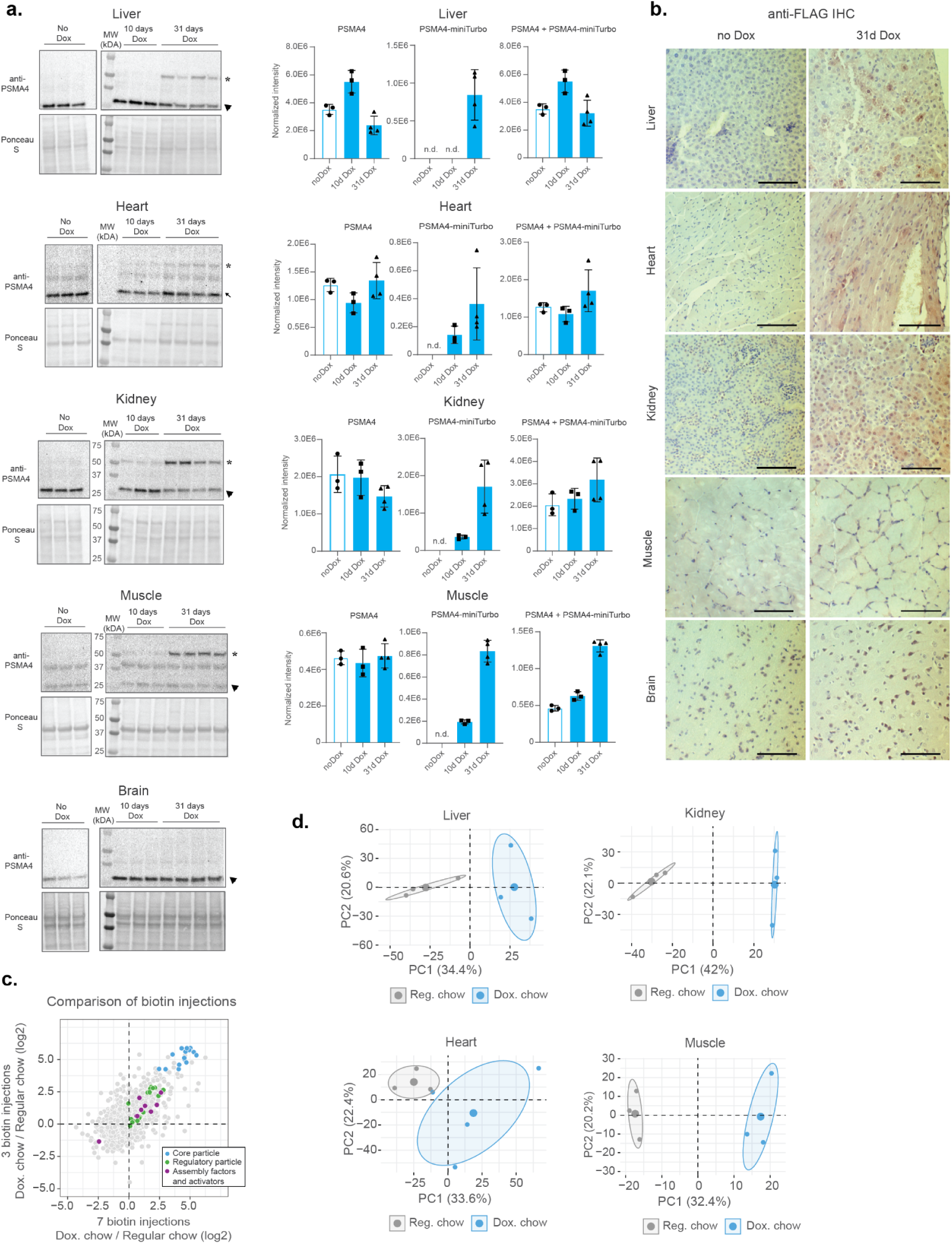
ProteasomeID mouse model validation and optimization. a. Immunoblots of PSMA4 protein performed on tissue lysates collected from ProteasomeID mice fed with regular chow or doxycycline containing food for 10 and 31 days and submitted to 3 daily biotin injections. Ponceau staining was used as loading control and for normalization of densitometric quantification of corresponding immunoblots. No Dox = mice fed with regular chow, 10d Dox = mice fed with doxycycline containing food for 10 days, 31d Dox = mice fed with doxycycline containing food for 31 days, error bars indicate standard deviation of the mean. b. Anti-FLAG immunohistochemistry analysis of tissues from ProteasomeID mice fed with regular chow (left) or doxycycline containing food (right) 31 days and submitted to 3 daily biotin injections. Scale bar indicates 20 µm. Dox = mice fed with doxycycline containing food; no Dox = mice fed with regular chow. c. Plot showing comparison of enrichment levels (in log2 fold change) of proteasome subunits and strong PIPs achieved in ProteasomeID mice fed with regular chow or doxycycline containing food for 10 and 14 days and submitted to 3 or 7 daily biotin injections respectively. d. Principal component analysis (PCA) of the tissue samples from ProteasomeID mice, based on the abundance of all proteins identified by label-free mass spectrometry. The smaller dots represent individual samples and the larger dots the centroids of each age-matched group. Ellipses represent 95% confidence intervals. The percentage of variance explained by the first two PC axes is reported in the axis titles.

The following tables are provided as Excel spreadsheets:

**Table S1:** BioID data from cell lines expressing different biotin ligase fusion proteins

**Table S2_Tab1:** SEC-MS data from HEK293T cells expressing PSMA4-BirA*

**Table S2_Tab2:** Biotinylation sites on proteasome subunits identified by PSMA4-BirA*

**Table S3_Tab1:** List of PIPs from Bousquet et al., Fabre et al., Gedaki et al.

**Table S3_Tab2:** List of PIPs from other studies

**Table S3_Tab3:** Protein groups enriched by ProteasomeID for PSMA4-BirA* dataset by the classifier algorithm and overlap with previous studies

**Table S3_Tab4:** GO enrichment candidate novel proteasome interactors

**Table S4_Tab1:** ProteasomeID data following treatment with MG132

**Table S4_Tab2:** Overlap between protein groups enriched only upon MG132 inhibition and the ones whose level of ubiquitination is shown to increase upon MG132 inhibition in a previous study

**Table S4_Tab3:** ProteasomeID data following treatment with PROTAC KB02-JQ1

**Table S5_Tab1-5:** Protein groups enriched by ProteasomeID for mouse dataset

**Table S5_Tab6:** Overlap of ProteasomeID enriched proteins between HEK293T and mouse organs

